# Current genomic deep learning models display decreased performance in cell type specific accessible regions

**DOI:** 10.1101/2024.07.05.602265

**Authors:** Pooja Kathail, Richard W. Shuai, Ryan Chung, Chun Jimmie Ye, Gabriel B. Loeb, Nilah M. Ioannidis

**Affiliations:** Center for Computational Biology, University of California, Berkeley, Berkeley, CA, USA; Department of Electrical Engineering and Computer Sciences, University of California, Berkeley, Berkeley, CA, USA; Division of Rheumatology, Department of Medicine, University of California, San Francisco, San Francisco, CA, USA; Institute for Human Genetics, University of California, San Francisco, San Francisco, CA, USA; Department of Epidemiology and Biostatistics, University of California, San Francisco, San Francisco, CA, USA; Bakar Computational Health Sciences Institute, University of California, San Francisco, San Francisco, CA, USA; Parker Institute for Cancer Immunotherapy, San Francisco, CA, USA; Chan Zuckerberg Biohub, San Francisco, CA, USA; Division of Nephrology, Department of Medicine, University of California, San Francisco, San Francisco, CA, USA; Cardiovascular Research Institute, University of California, San Francisco, San Francisco, CA, USA

**Keywords:** Deep Learning, Chromatin Accessibility, Variant Effect Prediction

## Abstract

**Background:** A number of deep learning models have been developed to predict epigenetic features such as chromatin accessibility from DNA sequence. Model evaluations commonly report performance genome-wide; however, *cis* regulatory elements (CREs), which play critical roles in gene regulation, make up only a small fraction of the genome. Furthermore, cell type specific CREs contain a large proportion of complex disease heritability.

**Results:** We evaluate genomic deep learning models in chromatin accessibility regions with varying degrees of cell type specificity. We assess two modeling directions in the field: general purpose models trained across thousands of outputs (cell types and epigenetic marks), and models tailored to specific tissues and tasks. We find that the accuracy of genomic deep learning models, including two state-of-the-art general purpose models – Enformer and Sei – varies across the genome and is reduced in cell type specific accessible regions. Using accessibility models trained on cell types from specific tissues, we find that increasing model capacity to learn cell type specific regulatory syntax – through single-task learning or high capacity multi-task models – can improve performance in cell type specific accessible regions. We also observe that improving reference sequence predictions does not consistently improve variant effect predictions, indicating that novel strategies are needed to improve performance on variants.

**Conclusions:** Our results provide a new perspective on the performance of genomic deep learning models, showing that performance varies across the genome and is particularly reduced in cell type specific accessible regions. We also identify strategies to maximize performance in cell type specific accessible regions.

## Background

Gene expression is regulated by nearby *cis* regulatory elements (CREs) such as promoters and enhancers. These CREs can be identified through functional epigenetic features such as chromatin accessibility, transcription factor (TF) binding, and histone marks. In the past several years, a number of deep learning models have aimed to predict and interpret these epigenetic features directly from DNA sequence [1–9]. A key application of these models is to probe the functional consequences of genetic variation within regulatory regions, particularly disease associated genetic variation. Current models have shown some promise in identifying causal variants within GWAS loci and annotating the mechanisms by which these variants act to modulate disease risk [1–4, 8]. Due to the wide variety of architectures and training procedures used by these models, a growing body of work seeks to perform systematic evaluations of the performance and limitations of genomic deep learning models for various tasks [10–15]. Genomic deep learning models are typically trained to maximize predictive performance on genome-wide assays. These genome-wide performance metrics place equal weight on all genomic regions, and may not be representative of performance within functional regulatory regions. CREs and, in particular, cell type specific CREs play critical roles in gene expression regulation and are known to harbor a large fraction of the heritability of complex diseases [16]. For this reason, we seek to further understand the advantages and limitations of current models with regard to their performance in regulatory regions with functional and disease relevance.

Here, we benchmark the performance of current genomic deep learning models in accessible regions with varying degrees of cell type specificity (Fig. 1A). We focus our analysis on predictions of chromatin accessibility, since accessibility characterizes potentially active CREs, and large chromatin accessibility datasets are publicly available for a diverse array of cell types, including at single-cell resolution, making it possible to robustly assess predictive accuracy in cell type specific accessible regions. We first study two recent state-of-the-art models: Enformer and Sei [7, 8] (Fig. 1B,C). Both Enformer and Sei are trained using multi-task learning over a large number of cell types and epigenetic marks, a common paradigm in the field. This approach was introduced by DeepSEA [1], which utilized 919 functional genomics tracks, and has been extended by Enformer and Sei to 5,313 and 21,907 tracks, respectively. Enformer predicts transcriptional activity, histone marks, TF binding, and chromatin accessibility and incorporates long-range sequence context up to 100kb away. Sei predicts histone marks, TF binding, and chromatin accessibility across more than 1,300 cell lines and tissues; to our knowledge, it is the genomic deep learning model trained on the largest set of chromatin accessibility profiles (2,372 profiles in total).

**Fig. 1.**
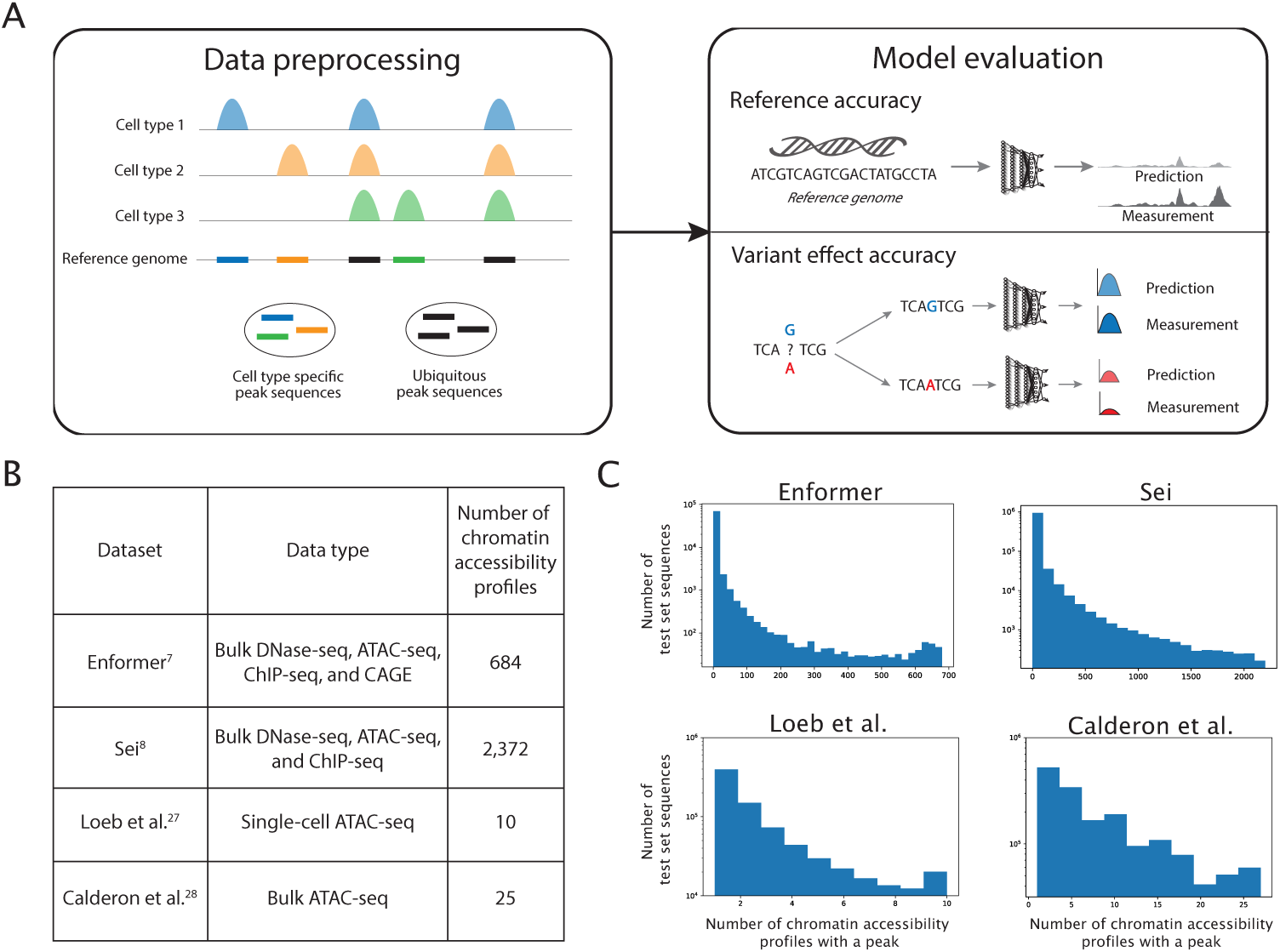
Overview of data processing and model evaluation. A) Schematic overview of the data preprocessing and evaluation pipeline used in this study. Cell type specific and ubiquitous peak sequences were annotated, and models were evaluated independently in these genomic regions. Models were evaluated on both “reference accuracy” (the models’ ability to predict experimentally measured accessibility from the reference genome) and “variant effect accuracy” (the models’ ability to predict allele-specific differences in accessibility). B) Four previously published datasets are used in subsequent analyses. The experimental assays and number of chromatin accessibility profiles are shown. Only chromatin accessibility profiles from ATAC-seq or DNase-seq are analyzed in this work. C) For each of the four datasets, the majority of test set sequences are cell type specific. Distributions shown are over test set sequences that had a peak in at least one chromatin accessibility profile in the dataset.

Although many genomic deep learning models are multi-tasked, no existing evaluations explore the impact of multi-tasking on predictions in cell type specific regions. In the absence of shared underlying features between tasks, it is possible for multi-tasking to decrease overall performance, a phenomenon known as negative transfer [17, 18]. Relatedly, even if cell types share regulatory grammar that may benefit from multi-task training, model capacity may be insufficient to learn the sequence features specific to each cell type as the number of cell types increases. Using custom models trained and evaluated on cell type specific ATAC-seq data from primary kidney and immune cells, we evaluate the effect of negative transfer and model capacity on predictive accuracy in cell type specific accessible regions by evaluating the performance of single-task and increased capacity multi-task models.

Another limitation of typical modeling assessments is that predictive performance is quantified by comparing experimental measurements to predictions made using the reference genome sequence. This type of evaluation – which we refer to as “reference accuracy” – does not directly measure a model’s ability to predict the effects of genetic variants. Using GWAS, eQTL, and allelic imbalance data, we evaluate variant effect predictions in accessible regions with varying degrees of cell type specificity.

Our evaluations provide insight into the performance of current state-of-the-art genomic deep learning models in accessible regions, and suggest strategies to maximize performance in cell type specific accessible regions.

## Results

### Evaluating state-of-the-art models in cell type specific accessible regions

Cell type specific accessible regions harbor much of the common genetic variation explaining heritability of human complex traits and diseases [16]. Therefore, we sought to characterize the ability of two state-of-the-art genomic deep learning models – Enformer and Sei – to predict chromatin accessibility in cell type specific accessible regions. Both models reportedly make highly accurate chromatin accessibility predictions as measured by the concordance between experimental and predicted accessibility across the whole genome. However, since cell type specific and ubiquitously accessible regions are regulated by different proteins and sequence determinants, model accuracy may differ in these regions.

We first verify the trait-relevance of cell type specific accessible regions in the Enformer training data, which includes 684 DNase-seq and ATAC-seq experiments from the ENCODE and Roadmap Epigenomics consortia [19, 20]. We categorize the 684 experiments (tracks) into 9 tissue categories, mirroring the categorization in [21], and divide the accessible regions (peaks) present in each tissue category into high and low cell type specificity subsets based on their overlap with peaks in the other accessibility tracks (Methods). For seven UK Biobank traits, we assess enrichment of trait heritability within these peak subsets using partitioned LD score regression [16]. We find that the high cell type specificity peaks from trait-associated tissues are highly enriched for trait heritability (Fig. 2A).

**Fig. 2.**
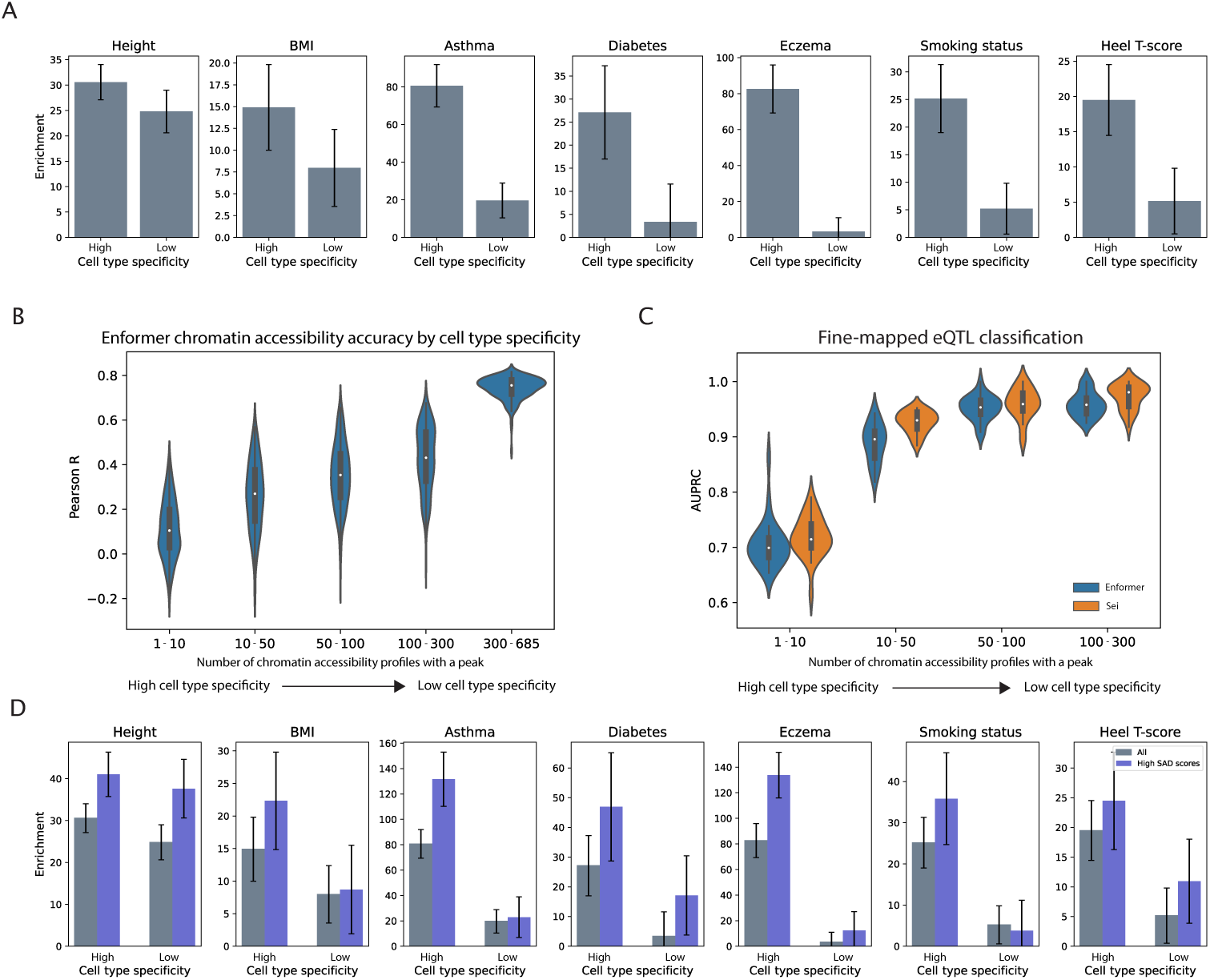
Evaluating state-of-the-art models in cell type specific peaks. A) Cell type specific peaks from trait-associated tissues represented in the Enformer training data are enriched for trait heritability. We categorize the 684 Enformer accessibility tracks into 9 tissue categories, mirroring the categorization in [21], and divide the accessible regions (peaks) present in each tissue category into high and low cell type specificity subsets based on their overlap with peaks in the other accessibility tracks (Methods). We compute heritability enrichments using the following trait-tissue associations – Height: Musculoskeletal-connective, BMI: Central nervous system, Asthma: Blood/immune, Dia-betes: Pancreas, Eczema: Blood/immune, Smoking status: Central nervous system, Heel T-score: Cardiovascular. B) Enformer’s chromatin accessibility prediction performance (reference accuracy) is poor in high cell type specificity peaks and highly accurate in low cell type specificity peaks (regions that contain a peak in greater than 300 chromatin accessibility profiles). Distributions shown are over all 684 Enformer accessibility output tracks. For the Sei model, which predicts the probability of the presence of a peak, we report the prediction AUC and AUPRC stratified by cell type specificity in Fig. S2. C) Enformer and Sei classify high posterior inclusion probability eQTLs (PIP *>* 0.9) versus a matched negative set of low PIP eQTLs (PIP *<* 0.01) (using positive and negative variant sets obtained from [7]). Both models have reduced performance when classifying eQTLs in cell type spe-cific accessibility peaks. D) Limited discrimination of trait-associated variants by Enformer variant effect predictions. Variants in chromatin accessible regions were subset to those with high Enformer SNP Accessibility Difference (SAD) scores (top 50% of Enformer SAD scores). Enrichment of these variants for trait heritability was assessed using partitioned LD score regression. We additionally report heritability enrichment for the top 10% of variants based on Enformer SAD scores in Fig. S6

To quantify the performance of Enformer and Sei in accessibility peaks with varying degrees of cell type specificity, we divide the test sequences for each model into bins based on the number of cell types in which that sequence has a peak in the experimental accessibility data. For the Enformer model, which predicts a continuous value corresponding to peak height, we report the Pearson correlation between the predicted and experimental accessibility for the test sequences in each bin (Fig. 2B), as well as the precision per peak for the cell types predicted to have the highest accessibility (Fig. S1). For the Sei model, which predicts a probability of the presence of a peak, we report the AUC and AUPRC for the predictions in each bin (Fig. S2). We observe that both Enformer and Sei make highly accurate predictions for sequences in the lowest cell type specificity bin (Enformer median Pearson R 0.76; Sei median AUC/AUPRC 0.99/0.99). However, the performance of both models drops on sequences that are cell type specific (Enformer median Pearson R 0.10; Sei median AUC/AUPRC 0.75/0.70 for the highest cell type specificity bin). To evaluate whether this drop in performance can be explained by lower experimental reproducibility for cell type specific peaks, we select five representative Enformer DNase tracks for which isogenic replicate data is available on ENCODE. For these tracks, we compare Enformer’s performance in each cell type specificity bin to the correlation in peak heights between isogenic replicates for the same set of peaks (Fig. S3). We observe only a slight decrease in isogenic replicate correlation for high cell type specificity peaks, which does not explain the dramatic drop in predictive accuracy.

Because the regulatory grammar at gene-proximal elements, such as promoters, is distinct from that at distal regulatory elements, we next ask whether reduced performance at cell type specific peaks is driven by distance between the peak and a gene transcription start site (TSS). We first evaluate the performance of Enformer for peaks stratified into three roughly equally sized TSS distance bins, and observe that while performance does decrease slightly for distal peaks (Fig. S4A, “All”), this decrease is minimal compared to the differences observed across the cell type specificity bins. Furthermore, stratifying by both TSS distance and cell type specificity demonstrates that cell type specificity and not TSS distance has a major impact on performance. Model predictions in high cell type specificity peaks are similarly poor regardless of whether the peaks are proximal or distal to a TSS (Fig. S4A). As a higher fraction of cell type specific peaks are distal to a TSS (Fig. S4B), cell type specificity may contribute to the decreased performance at distal peaks.

Having established that Enformer and Sei’s reference accuracy is lowest in peaks with the highest degree of cell type specificity, we next evaluate whether this trend in performance extends to the models’ ability to predict the functional effects of single nucleotide polymorphisms (SNPs), or “variant effect accuracy”. We perform two evaluations to assess variant effect accuracy, using both eQTLs and GWAS heritability enrichment.

First, we utilize fine-mapped eQTLs from the Genotype-Tissue Expression (GTEx) Consortium [22]. We divide the positive set of fine-mapped GTEx eQTLs with high posterior inclusion probability (PIP *>* 0.9) into bins based on the cell type specificity of the accessibility peak they overlap. Using variant effect predictions for all accessibility tracks from either Enformer or Sei, we train random forests to classify the positive set eQTLs in each bin versus a matched negative set of low PIP eQTLS (PIP *<* 0.01) (Fig. 2C), following a methodology similar to [7]. We use only chromatin accessibility variant effect predictions from Enformer and Sei in these evaluations, since our goal is to understand how chromatin accessibility predictions vary in regions with varying degrees of cell type specificity. We observe that both Enformer and Sei perform best at classifying eQTLs in the low cell type specificity bin (Enformer median AUPRC 0.93; Sei median AUPRC 0.96), with decreasing performance in high cell type specificity bins (Enformer median AUPRC 0.71; Sei median AUPRC 0.71 in the highest cell type specificity bin). These results suggest that variant effect prediction remains more challenging within cell type specific peaks.

In a similar manner, we test each model’s ability to predict the direction of effect – the eQTL sign – of high PIP eQTLs in accessible regions with varying degrees of cell type specificity (Fig. S5). Avsec et al. [7] showed that Enformer’s variant effect predictions are somewhat predictive for this task, but much less than for classifying high vs. low PIP eQTLs. As with the previous classification task in Fig. 2C, we use only chromatin accessibility variant effect predictions in this evaluation. We observe that both Enformer and Sei perform most poorly on direction-of-effect prediction in the high cell type specificity bin (Enformer median AUC 0.59; Sei median AUC 0.58), with improving performance in low cell type specificity bins (Enformer median AUC 0.77; Sei median AUC 0.73 in the lowest cell type specificity bin).

Overall, we find that Enformer and Sei’s ability to predict variant effects on gene expression decreases for variants within cell type specific accessibility peaks. We consider the possibility that these differences in performance could be explained by differences in distance to the TSS or effect size of the eQTLs. We observe only small differences in TSS distance or effect size distributions for eQTLs in high or low cell type specificity bins (Fig. S5C,D). Therefore, it is unlikely that systematic differences in effect size or TSS distance entirely explain the differences we observe in the ability of Enformer and Sei to predict cell type specific eQTLs.

As a second assessment of variant effect accuracy, we test whether variants with larger predicted differences in accessibility are enriched for trait heritability within each of the high and low cell type specificity peak subsets from the heritability analysis in Fig. 2A. We subset the variants in these peaks into two groups – those with low versus high absolute SNP accessibility difference (SAD) scores – based on Enformer’s variant effect predictions. Using partitioned LD score regression, we assess enrichment of trait heritability among the variants with high Enformer SAD scores. For the majority of tested traits, we find that variants with high (top 50%) Enformer SAD scores in cell type specific peaks are more enriched for trait heritability than all variants in cell type specific peaks (Fig. 2D). For some traits – including Heel T-score – Enformer’s SAD scores are less effectively able to identify the trait-relevant variants within cell type specific peaks (Fig. 2D). Subsetting the variants in low cell type specificity peaks to those with high Enformer SAD scores is less informative for identifying trait-relevant genetic variation, but this is likely due to the fact that the low cell type specificity peaks are less enriched for trait heritability to begin with. To verify that the above findings are robust to the threshold used to define high SAD score variants, we also analyze the subset of variants with the top 10% highest Enformer SAD scores (Fig. S6).

Taken together, these results indicate that state-of-the-art models have decreased reference accuracy in cell type specific accessible regions, and their ability to make accurate variant effect predictions within these regions may depend on the specific task or phenotype of interest.

### Multi-task models trained on related cell types exhibit poor cell type specific accessibility prediction

In addition to evaluating state-of-the-art genomic deep learning models trained on large compendia of data from diverse cell types, we also consider an alternate direction in the field – training bespoke deep learning models on smaller datasets to interrogate a specific biological system or disease [23–26]. Unlike Enformer and Sei – which are tasked with learning the regulatory grammar of a wide array of cell types – these bespoke models are usually trained on a small number of related cell types, which may share more regulatory logic. We reasoned that this framework might be more amenable to learning cell type specific regulatory syntax, and sought to also benchmark the performance of such bespoke models in cell type specific accessible regions.

To this end, we utilize two ATAC-seq datasets – single-cell ATAC-seq of primary human kidney tissue from three donors [27] and bulk ATAC-seq of 25 primary human immune cell types, sorted by flow cytometry, from four human blood donors [28] (Fig. 1B,C). In each dataset, accessibility peaks were grouped into disjoint clusters based on their accessibility profiles across cell types. This clustering results in one cluster for each dataset corresponding to ubiquitously accessible peaks and additional clusters displaying cell type specificity. To verify the disease-relevance of the cell type specific peaks in both ATAC-seq datasets, we estimate the enrichment of trait heritability in these ubiquitous or cell type specific accessibility clusters using partitioned LD score regression for the kidney function marker creatinine (for the Loeb et al. [27] data) or immune-related traits (for the Calderon et al. [28] data) in the UK Biobank. In the Loeb et al. [27] data, we find that tubule cell type specific peak clusters – accessible specifically in proximal tubule, distal tubule, or loop of Henle cells – are significantly enriched for creatinine heritability (18-fold enrichment in proximal tubule specific peaks, 15-fold enrichment in distal tubule/loop of Henle specific peaks) (Fig. 3A), similar to the enrichment of ubiquitous peaks. In the Calderon et al. [28] data, we find that cell type specific peak clusters are significantly enriched for heritability of the immune-related traits asthma and eczema (14-fold average enrichment in myeloid specific peaks, 29-fold average enrichment in NK cell specific peaks, and 49-fold average enrichment in T cell specific peaks) (Fig. 3A). In ubiquitous peaks, we observe no evidence of significant immune-related trait heritability enrichment.

**Fig. 3.**
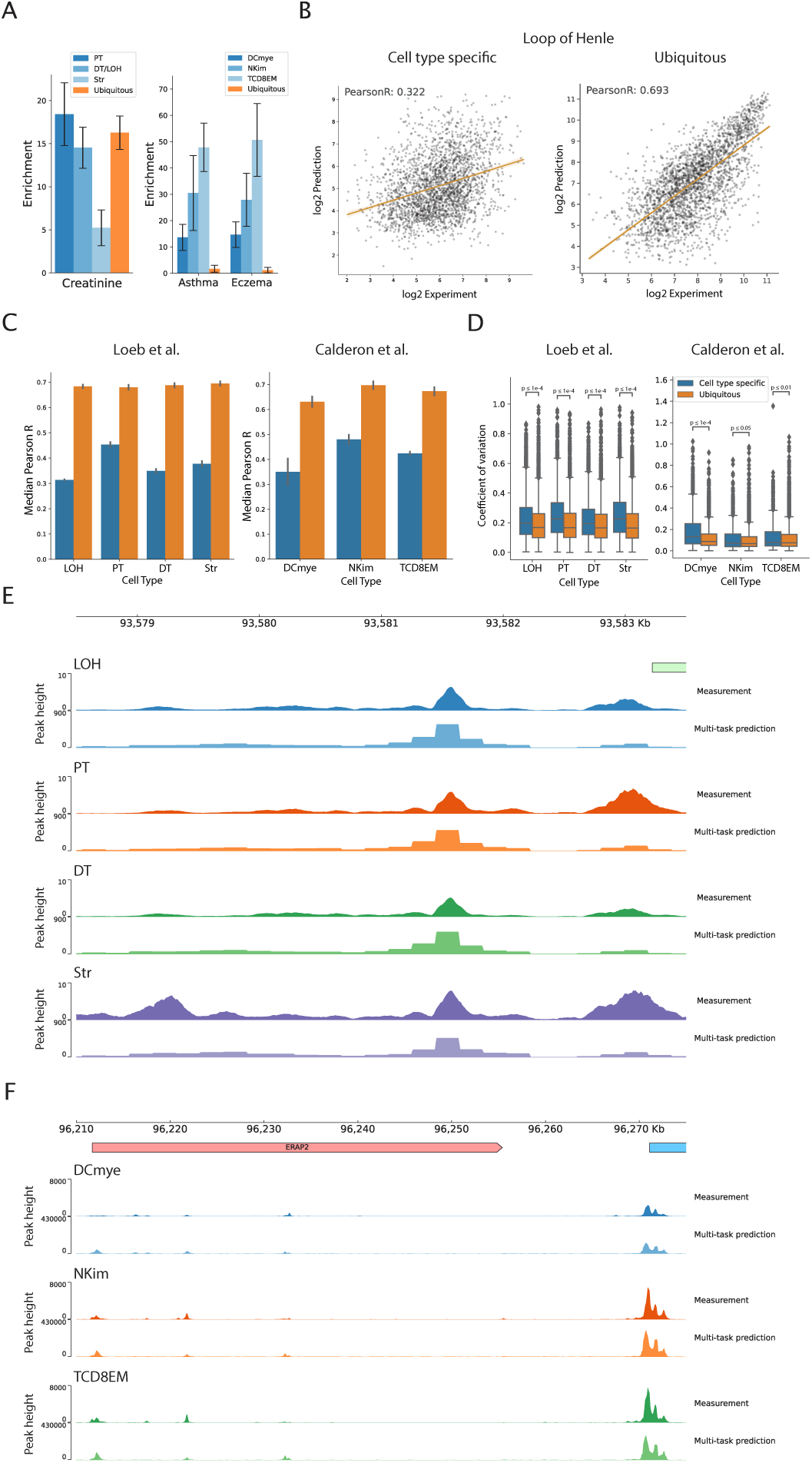
Multi-task accessibility prediction models of related cell types exhibit poor cell type specific peak prediction. A) Kidney tubule cell type specific accessibility peaks are signif-icantly enriched for heritability of the kidney function biomarker creatinine (Loeb et al. [27] data) and immune cell type specific accessibility peaks are significantly enriched for autoimmune trait her-itability (Calderon et al. [28] data). B) Scatter plots of experimentally measured versus predicted accessibility in cell type specific and ubiquitous peaks for one cell type – Loop of Henle – in the Loeb et al. [27] data. Plotted points are sequences from the held out test chromosomes. C) Multi-task model reference accuracy is poor in cell type specific peaks for multi-task models trained on either the Loeb et al. [27] data or the Calderon et al. [28] data. Reference accuracy is measured as the Pear-son correlation between experimentally measured versus predicted accessibility. Error bars represent the standard deviation over three replicate models. D) Multi-task model predictions across repli-cate models are significantly more variable for sequences in cell type specific peaks versus sequences in ubiquitous peaks. Variability is quantified as the coefficient of variation for each sequence across three model replicates (one-sided Mann-Whitney U-test with Benjamini-Hochberg multiple testing correction). E) Experimentally measured and predicted accessibility profiles from the Loeb et al. [27] data for a region around *NR2F1*. The ubiquitous peak near the center of the coverage track is well-predicted in all cell types by the multi-task model, while the cell type specific peak on the 5’ end of the coverage track is not predicted to be a peak in any cell type. F) Experimentally measured and predicted accessibility profiles from the Calderon et al. [28] data for a region around *ERAP2*. The ubiquitous peak on the 3’ end of the coverage track is well-predicted in all cell types by the multi-task model. The two cell type specific peaks towards the 5’ end of the coverage track are predicted to be peaks in all three cell types by the same model, although there is no measured accessibility in these regions in DCmye cells.

For each dataset, we then train a set of multi-task convolutional neural networks (CNNs) to map input DNA sequences (1344bp) to continuous measures of chromatin accessibility (normalized ATAC-seq read counts) in each cell type. Our architecture is based on an updated version of the Basset model [2]. We train three replicates of each model to assess uncertainty in model predictions. All models achieve a reference accuracy on held out test chromosome sequences (Table S1, Table S2) that is comparable to the results reported in previous work [3]. Quantifying the predictive performance of these multi-task models in cell type specific and ubiquitous peaks separately, we observe that performance in cell type specific peaks (0.39 avg. Pearson R for Loeb et al. [27]; 0.30 for Calderon et al. [28]) is markedly lower than in ubiquitous peaks (0.69 avg. Pearson R for Loeb et al. [27]; 0.68 for Calderon et al. [28]) (Fig. 3B,C). As a measure of uncertainty in model predictions, we compute the coefficient of variation for each sequence across the three trained model replicates. We find that predictions across the different models are significantly more variable for sequences in cell type specific peaks than sequences in ubiquitous peaks (Fig. 3D). These results indicate that deep learning models trained using multi-task learning on related cell types also have decreased performance and greater uncertainty within cell type specific peaks.

We next investigate whether various characteristics of cell type specific peaks might explain some of the differences in model performance. We first test for systematic differences in the degree of accessibility between cell type specific and ubiquitous peaks by quantifying the distribution of peak heights, since many ubiquitous peaks are at promoters and many cell type specific peaks are at distal enhancers. We find that cell type specific peaks tend to have lower peak heights than ubiquitous peaks in the single-cell Loeb et al. [27] data (Fig. S7A), but we do not observe a similar trend in the bulk Calderon et al. [28] data (Fig. S7B). After controlling for this bias in the Loeb et al. [27] data, we still observe a drop in performance in cell type specific peaks when compared to height-matched ubiquitous peaks (0.39 vs 0.50 avg. Pearson R) (Fig. S7C).

We also investigate whether ubiquitous peaks have more easily recognizable sequence features than cell type specific peaks, which might make them easier for a model to learn. We find that ubiquitous peaks have slightly higher GC content than non-ubiquitous peaks (Fig. S8A) and are more likely to contain putative CpG islands and CTCF motifs (Fig. S8B,C). We perform a motif enrichment analysis for ubiquitous and cell type specific peaks and identify several TFs previously known to be active in the studied cell types (Fig. S8D); for example, we identify HNF4A as enriched in proximal tubule peaks, SPI1 as enriched in myeloid cell peaks, and Jun as enriched in T cell peaks. Particularly in the Loeb et al. [27] data, we find stronger enrichment of motifs in ubiquitous peaks compared to cell type specific peaks (Fig. S8D).

To illustrate model performance at cell type specific peaks, we present example loci from each dataset (Fig. 3E,F). At the *NR2F1* locus, the multi-task model trained on the Loeb et al. [27] data accurately predicts accessibility at a ubiquitously accessible peak, but fails to identify a peak in any cell type at a nearby cell type specific peak (Fig. 3E). Similarly, at the *ERAP2* locus, the multi-task model trained on the Calderon et al. [28] data accurately predicts accessibility at a ubiquitously accessible peak, but predicts a small amount of accessibility in all cell types at the cell type specific peaks nearby (Fig. 3F).

### Increased capacity to learn cell type specific regulatory syntax improves cell type specific accessibility prediction

To provide insights to help guide future modeling improvements, we next characterize the effect of a number of common training decisions on cell type specific accessibility prediction (Fig. 4A; Fig. S9A). First, a multi-task architecture might cause models to learn shared rather than cell type specific features, leading to higher performance in ubiquitous peaks than in cell type specific peaks. Since learning shared features could cause over-correlated predictions across cell types, we compare correlations across cell types for both predicted and experimentally measured peak heights. As expected, cell type specific peaks exhibit low correlation in measured accessibility across cell types (Fig. 4B, in gray). However, the multi-task model’s predictions in cell type specific regions are highly correlated between cell types (Fig. 4B, in dark blue). Ubiquitously accessible peaks exhibit high correlation in measured accessibility across cell types, with modest over-correlation in predicted accessibility across cell types (Fig. S9B).

**Fig. 4.**
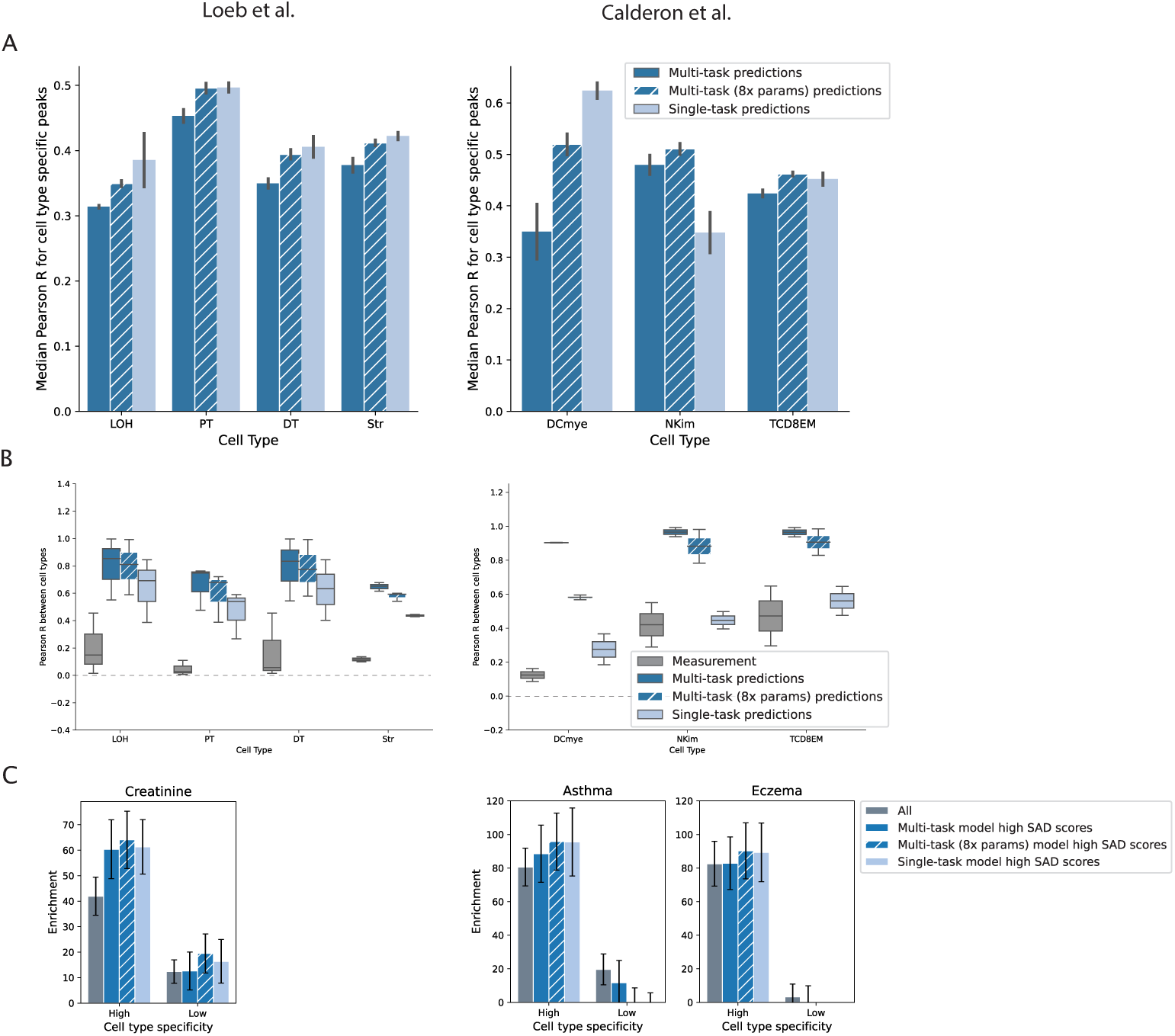
Increased model capacity to learn cell type specific regulatory syntax improves reference sequence prediction in cell type specific peaks. A) Reference accuracy of multi-task versus single-task models evaluated in cell type specific peak regions. Single-task models and high capacity multi-task models tend to outperform baseline multi-task models in cell type specific peaks. Reference accuracy of the same multi-task and single-task models in ubiquitous peaks is reported in Fig. S9A. B) In cell type specific peaks, pairwise correlations of peak height between cell types are computed for experimental (dark gray) and model predicted accessibility (dark and light blue). Model predicted accessibility is more correlated between cell types than experimentally measured accessibility, and this overcorrelation is more pronounced in predictions from multi-task models than predictions from single-task models. The correlation in experimental and model predicted accessibility between cell types in ubiquitous peaks is reported in Fig. S9B. C) High SAD score variants from all three tested model types (multi-task, high capacity multi-task, and single-task) are similarly enriched for trait heritability of tissue-matched traits. Using predictions from each model, we subset the variants in high and low cell type specificity peak regions based on the model’s SNP Accessibility Difference (SAD) scores. We use the median SAD score for all variants in a particular peak set (e.g. “Kidney high cell type specificity peaks”) as a threshold to subset to high SAD score variants. Enformer’s performance on this task for the same traits is shown in Fig. S19.

To determine whether the observed over-correlation in predicted versus experimentally measured accessibility between cell types can be attributed to experimental measurement noise, we also measure the correlation across individuals of experimental accessibility in the same cell type (Fig. S10). We observe high correlation between individuals within cell type specific peaks (Fig. S10A, in light gray; Loeb et al. [27] median Pearson R 0.72; Calderon et al. [28] median Pearson R 0.72), which accounts for both biological and experimental sources of variation. Thus, experimental noise does not explain the low correlation of measured accessibility between cell types at cell type specific peaks. We conclude that over-correlation between predictions for different cell types, which is most pronounced within cell type specific peaks, is a characteristic of these multi-task models.

We then train single-task models on each cell type individually, to test whether this over-correlation is caused by the multi-task architecture. Single-task training yields a small drop in genome-wide test set performance (Fig. S11A, Fig. S12A), but leads to a performance improvement in cell type specific peaks (Fig. 4A). We also evaluate whether a transfer learning approach, in which a single-task model is first trained on a cell type with abundant high-quality data, and then fine-tuned on a cell type with lower quality data, improves performance for cell types with lower quality data (reasoning that this approach might retain beneficial aspects of both multi-task and single-task models). However, this transfer learning approach leads to worse performance in cell type specific regions than single-task training, even using a relatively sparse dataset (Fig. S13). For this reason, we primarily compare single-task and multi-task models in subsequent evaluations. We find that single-task training reduces the spurious over-correlation between cell types observed for the multi-task models (Fig. 4B, Fig. S9B, in lighter shade), and generally reduces false positive peak predictions (Fig. S14).

The performance improvements that we observe with single-task learning could be due to minimizing the effect of negative transfer across cell types, or due to increased capacity to learn cell type specific regulatory syntax. Therefore, we also train additional multi-task models with increased capacity (2, 4, and 8 times the number of parameters) compared to the original baseline multi-task model. We find that increasing model capacity has little to no effect on performance genome-wide (Fig. S15A,B, in gray) or in ubiquitous peaks (Fig. S9A,B, in hatched orange; Fig. S15A,B, in orange), but does improve performance in cell type specific peaks (Fig. 4A,B, in hatched blue; Fig. S15A,B, in blue). In most cases, the performance of the highest capacity multi-task model is comparable to single-task performance, suggesting that the performance improvements from single-task learning are primarily due to increased capacity to learn cell type specific regulatory syntax.

Differences in cell type specific prediction between models are likely to be driven by differences in their ability to recognize cell type specific TF motifs, and to correctly associate these motifs with accessibility in particular cell types. To assess this, we first perform a motif insertion analysis using the baseline multi-task, high-capacity multi-task, and single-task models (Methods). In both the Loeb et al. [27] and Calderon et al. [28] datasets, we observe greater heterogeneity between cell types in model-predicted TF activity for the single-task models as compared to the baseline multi-task model (Fig. S16A,B). For example, in the Loeb et al. [27] data, the single-task models – as well as, to a lesser extent, the high-capacity multi-task model – identify SRF activity as restricted to Stromal cells, consistent with previous reports [27] (Fig. S16A). The baseline multi-task model fails to learn this TF as active in any cell type. In the Calderon et al. [28] data, the single-task models and high-capacity multi-task model identify SPI1 as a myeloid-specific TF, while the baseline multi-task model instead predicts only weak SPI1 activity across cell types (Fig. S16B). Similarly, the single-task models predict more specific TF activity for many FOS/JUN TFs in CD8 T cells, while the baseline multi-task model predicts these TFs to be broadly active. Both SPI1 and FOS/JUN TFs have previously been characterized to have cell type specific activity in myeloid and T cells, respectively [28]. Second, we use TF-MoDISco [29] to identify the TF motifs that drive predictions in ubiquitous and cell type specific peaks for the baseline multi-task, high-capacity multi-task, and single-task models.

We compute the Jaccard similarity in the motifs identified by TF-MoDISco across cell types, and find that the motifs driving single-task and high-capacity multi-task model predictions tend to be more distinct across cell types, particularly in cell type specific peaks (Fig. S16C,D). Taken together, these observations suggest that some of the over-correlation between cell types that we observe for the baseline multi-task models may be due to the models learning a similar regulatory syntax for distinct cell types. However, we note that even single-task models still suffer from over-correlated predictions between cell types (Fig. 4B; Fig. S9B), suggesting that there is still a bias towards learning shared sequence features that is architecture agnostic.

For two example cell type specific peaks, we examine the sequence features that contribute to distinct predictions between different model types. In the Loeb et al. [27] data, we highlight a Stromal cell specific peak that is mispredicted by the baseline multi-task model to be a peak in additional cell types, but is correctly predicted by the single-task models to only be accessible in Stromal cells. *In silico* mutagenesis (ISM) of this peak reveals that a TEAD-like motif is contributing to the predictions of the baseline multi-task model in both Stromal and non-Stromal cell types, while this TEAD-like motif has less importance in non-Stromal cell types for the high-capacity multi-task and single-task models (Fig. S17A). Some of the difference in model predictions may also be due to differences in the scale of ISM scores observed across the entire region for different models. (Note that we compare ISM scores on the same scale for all models, since we generally observe that the models make predictions on similar scales for held-out reference sequences.) Similarly, in the Calderon et al. [28] data, we highlight a dendritic cell specific peak that is mispredicted by the baseline multi-task model to be a peak in additional cell types, but is correctly predicted by the single-task models to only be accessible in dendritic cells. ISM of this peak reveals that a SPIB-like motif is weakly informing predictions of the baseline multi-task model in dendritic and non-dendritic cell types, while this SPIB-like motif is more strongly and specifically driving dendritic cell predictions for the high-capacity multi-task and single-task models (Fig. S17B). Note that this pattern mirrors the differences in predicted TF activity for SPIB in the motif insertion analysis above (Fig. S16B), where we observe stronger and more specific predicted SPIB activity in dendritic cells with high-capacity multi-task and single-task models.

We also examine predictions from the high capacity multi-task and single-task models for the *NR2F1* and *ERAP2* loci where we previously observed incorrect cell type specific predictions from the baseline multi-task models (Fig. 3E,F). We find that for the Stromal cell specific peak at the *NR2F1* locus, both the high capacity multi-task model and single-task models still fail to accurately capture its peak height in Stromal cells (Fig. S18A). For the cell type specific peaks at the *ERAP2* locus, the high capacity multi-task and single-task models reduce false positive predictions at some cell type specific peaks, but continue to demonstrate limited precision as well as some false negative predictions at other cell type specific peaks (Fig. S18B). Taken together, our observations in these and the above example loci reflect the general trend of modest improvement in cell type specific peak reference accuracy from increasing model capacity to learn cell type specific motifs, either through single-task learning or high-capacity multi-task models.

We next sought to evaluate whether increasing model capacity also affects variant effect accuracy, using two types of evaluations. First, we use allelic imbalance measurements at heterozygous sites in the experimental ATAC-seq data to test the models’ ability to predict the higher accessibility allele. In both datasets, we do not observe consistent improvements in chromatin accessibility allelic imbalance prediction by single-task models (Fig. S11B, Fig. S12B). Second, we compare the models on the GWAS heritability benchmark, paralleling the analysis in Fig. 2D, using tissue-matched traits (Creatinine for the Loeb et al. [27] data; Asthma and Eczema for the Calderon et al. [28] data). We find that high SAD score variants from single-task, high-capacity multi-task, and baseline multi-task models are all similarly enriched for trait heritability (Fig. 4C). Together, these results indicate that the higher capacity models tested here do not substantially improve variant effect prediction, even in cell type specific accessible regions. For reference, we also compare the heritability enrichments obtained with these tailored tissue-specific models to those obtained with Enformer for matched tissue tracks (Fig. S19). For Creatinine, high SAD score variants from the tissue-specific models have similar enrichment to high SAD score variants from Enformer (Fig. S19A). However, for the immune-related traits, Enformer’s high SAD score variants are significantly more enriched for trait heritability than high SAD score variants from the tissue-specific models (Fig. S19B). Differences in performance may be due to the higher capacity model architecture of Enformer versus the smaller Basset-style architecture of the tissue-specific models tested here, or to Enformer’s training on a larger number of cell types, tissues, and assays (including CAGE-seq and ChIP-seq). One possible explanation for the difference between immune and kidney traits in the relative performance of Enformer is the makeup of different track types in the Enformer training data. In particular, immune cell types and tissue samples are heavily overrepresented, which may bias Enformer towards learning immune cell regulatory syntax. For other cell types, such as kidney, variant effect predictions from significantly smaller models trained on limited data from the relevant cell types can perform similarly to Enformer.

### Choice of training regions does not substantially impact cell type specific accessibility prediction

The second modeling choice that we explore is which regions of the genome to include in training. For the kidney and immune-specific single-task and multi-task models described above, we include all regions of the genome from the training set chromosomes, apart from assembly gaps and unmappable regions. Cell type specific accessible regions make up less than 10% of this training set. To test whether the relative infrequency of these sequences in the training set contributes to poor performance, we evaluate five different training sets with varying proportions of cell type specific accessible regions (Methods, Table 1). For each training set, we train models using both multi-task and single-task learning, and evaluate their performance in both cell type specific and ubiquitously accessible regions (Fig. S11A,B, Fig. S12A,B). In addition to evaluating the reference accuracy of each training decision, we also evaluate variant effect accuracy using the chromatin accessibility allelic imbalance data. We observe that for most choices of training set, single-task learning improves reference accuracy in cell type specific accessible regions and has minimal effect in ubiquitously accessible regions. The choice of training set yields different results for the two datasets. For the Loeb et al. [27] kidney data, the training set with the highest proportion of cell type specific accessible sequences yields the highest reference accuracy in cell type specific accessible regions. However, for the Calderon et al. [28] immune data, the training set that includes all genomic sequences yields the highest reference accuracy in cell type specific accessible regions. For variant effect accuracy, we do not observe consistent trends for the training sets we evaluated, but we note that training on all genomic sequences performs as well or better than the other training choices in almost all cases. Although our ability to measure variant effect accuracy using allelic imbalance is limited by the small number of individuals assayed, and thus the number of heterozygous sites, based on these data we do not observe that a particular choice of training set consistently improves performance in cell type specific accessible regions.

**Table 1.** Description of evaluated training sets.

| Training set | Genomic regions |
| --- | --- |
| 1 | All genomic sequences |
| 2 | Sequences overlapping any ATAC peak and an equal number of non-peak sequences (1:1 peak:non-peak ratio) |
| 3 | Sequences overlapping any ATAC peak and an equal number of GC-matched non-peak sequences (1:1 peak:non-peak ratio) |
| 4 | Sequences overlapping any ATAC peak |
| 5 | Sequences overlapping non-ubiquitous ATAC peaks |

## Discussion

We have performed a systematic analysis of genomic deep learning models that predict chromatin accessibility from DNA sequence, focusing on accessible regions of the genome with varying degrees of cell type specificity. While most previous evaluations of genomic deep learning models have focused on genome-wide performance metrics, which may mask performance differences on small but biologically important subsets of the genome, here we evaluated performance independently in different genomic regions. We found that predictive performance varies dramatically across the genome, and is particularly poor in cell type specific accessible regions, which are known to harbor a large fraction of disease heritability. This finding is consistent both for general purpose models such as Enformer and Sei, which are trained on large compendia of publicly available data, as well as models trained on smaller tissue-specific datasets. We performed additional variant-based evaluations using eQTL and GWAS data and found that eQTL variant effect prediction accuracy also decreases in cell type specific accessible regions. Previous work has demonstrated that genomic deep learning models perform more poorly on distal eQTLs [7, 12]. Our results demonstrate that much of this effect may be explained by the increased cell type specificity of the regulatory elements harboring these distal eQTLs.

We also highlight the importance of the choice of performance metric in model evaluations. The performance metric commonly used to evaluate genomic deep learning models is the concordance between experimental measurements and model predictions for input sequences from the reference genome; this “reference accuracy” metric does not directly measure a model’s ability to predict variant effects. Using multiple model types and training data sets, we observed that models with improved performance to predict chromatin accessibility using the reference genome often do not demonstrate improved performance in variant effect prediction tasks. As many of the most important applications of genomic deep learning models are for variant effect prediction, these results, as well as a growing body of literature [10, 12–14], imply that an important direction for the field is to comprehensively evaluate models on their utility for variant interpretation tasks, independently of reference sequence performance.

Finally, we characterized the effect of a number of common training decisions on cell type specific accessibility prediction to provide insight that may help guide future modeling improvements. In one previous study, Maslova et al. [23] evaluated the choice of loss function on cell type specific accessibility prediction. They found that using a Pearson correlation loss function, which directly emphasizes accessibility differences between cell types, improves cross-cell type predictions in cell type specific peaks, as compared to a mean-squared error loss function. In this work, we evaluated models trained using multi-task and single-task learning, as well as different choices for training set composition. We found that when compared to baseline multi-task models, single-task models and higher capacity multi-task models improved reference sequence prediction in cell type specific accessible regions. These results highlight the importance of careful evaluation on biologically relevant genomic regions and tasks in designing model architectures, as higher capacity models did not provide any meaningful improvement on standard genome-wide reference accuracy metrics.

## Conclusions

In summary, we demonstrated that the performance of current genomic deep learning models varies dramatically across the genome and is particularly poor in cell type specific accessible regions, which harbor a large fraction of the heritability of human diseases. We characterized the effects of a number of common training decisions on cell type specific accessibility prediction, and identified single-task learning and high capacity multi-task models as potential methods to improve reference sequence prediction in cell type specific accessible regions. Overall, these evaluations provide a new perspective on the performance of current genomic deep learning models, and suggest paths to maximize performance in cell type specific accessible regions.

## Declarations

### Ethics approval and consent to participate

Not applicable.

### Consent for publication

Not applicable.

### Availability of data and materials

**Enformer model and data:** The pre-trained Enformer model was obtained from https://tfhub.dev/deepmind/enformer/1. Enformer training, validation, and test data was obtained from https://console.cloud.google.com/storage/browser/basenji barnyard/data. Pre-computed variant effect predictions for all frequent variants in the 1000 Genomes cohort were obtained from https://console.cloud.google.com/storage/ browser/dm-enformer/variant-scores/1000-genomes/enformer.

**Sei model and data:** The pre-trained Sei model (https://zenodo.org/records/ 4906997) and relevant resources (https://zenodo.org/records/4906962) were obtained from Zenodo [30, 31]. Sei test data and predictions were obtained from S3 using instructions provided in the Sei manuscript Github repository (https://github.com/ FunctionLab/sei-manuscript).

**Loeb et al.** [27] **data:** Processed ATAC-seq data from [27] were obtained from GEO accession GSE262931.

**Calderon et al.** [28] **data:** Bigwig files used to train models were obtained from https://s3.amazonaws.com/muellerf/data/trackhubs/immune atlas/hg19/. Cell type specific peak clusters and allelic imbalance data were obtained from Supplementary Table 1 of [28].

**Models trained on Loeb et al.** [27] **and Calderon et al.** [28] **data**: Model weights for models trained on the Loeb et al. [27] and Calderon et al. [28] data can be downloaded from Zenodo (https://zenodo.org/records/10729956) [32].

**Additional datasets:** GTEx SuSiE fine-mapped eQTL data and matched negative sets were obtained from https://console.cloud.google.com/storage/ browser/dm-enformer/data/gtex fine. UK Biobank GWAS summary statistics were obtained from the Neale lab server using the following script https://github.com/ni-lab/CellTypeSpecificAccessibilityPrediction/blob/main/ scripts/enformer/ldsc/download gwas sumstats.sh. TF binding profiles for the motif enrichment and model interpretability analyses were obtained from the JASPAR 2022 CORE vertebrates non-redundant collection [33]. Isogenic replicate data for selected Enformer tracks were obtained from the ENCODE portal [34] (https://www.encodeproject.org/) with the following identifiers: ENCFF102UQK (track 79), ENCFF492TUE (track 79 replicate), ENCFF634ZUJ (track 96), ENCFF089MHS (track 96 replicate), ENCFF457RRO (track 112), ENCFF064VXK (track 112 replicate), ENCFF302JEV (track 135), ENCFF241ZSS (track 135 replicate), ENCFF827VFY (track 144), ENCFF524NIB (track 144 replicate).

**Code:** Code used in the current study, as well as additional instructions to download datasets, are available in the Github repository (https://github.com/ni-lab/ CellTypeSpecificAccessibilityPrediction) and on Zenodo (10.5281/zenodo.11588989) [35].

### Competing interests

The authors declare that they have no competing interests.

### Funding

This work was partially supported by the U.S. National Institutes of Health grants R00HG009677 and R01HG011239, funding from the UC Berkeley-UCSF Faculty Collaboration Program, and grants from the UC Noyce Initiative for Computational Transformation and Chan Zuckerberg Initiative. C.J.Y. is an investigator at the Chan Zuckerberg Biohub and is a member of the Parker Institute for Cancer Immunotherapy (PICI). N.M.I. is a Chan Zuckerberg Biohub Investigator.

### Author contributions

P.K., G.B.L., and N.M.I. conceived the study. P.K., R.W.S., and R.C. analyzed the data. All authors analyzed the results. P.K., G.B.L., and N.M.I. wrote the manuscript with input from all authors.

## Acknowledgements

We thank David Kelley, Xinming Tu and members of the Ioannidis and Ye labs for helpful discussions.

## Additional files

**Supplementary Tables:** Table S1: Loeb et al. [27] multi-task model performance. Table S2: Calderon et al. [28] multi-task model performance.

## Methods

### Chromatin accessibility datasets

Four chromatin accessibility datasets were used throughout this study. Briefly, we describe each of these datasets and the additional data processing steps we performed.

### Enformer data

We obtained processed training, validation, and test data used to train Enformer from [7]. These data contained 684 chromatin accessibility profiles from the ENCODE [19] and Roadmap Epigenomics [20] consortia that had been processed in [36] to summarize the read coverage for each profile in 128-bp bins along the genome. For each 128-bp bin in the test data, we called peaks on the read coverage values from each chromatin accessibility profile using a Poisson model parameterized by a global null lambda similar to the MACS2 approach [37], and applied a 0.01 FDR cutoff. All data were processed using the hg38 reference genome.

### Sei data

We obtained processed training, validation, and test data used to train Sei from [8]. These data contained 2,372 chromatin accessibility profiles from the Cistrome, ENCODE, and Roadmap Epigenomics consortia [19, 20, 38]. In contrast to the Enformer data, which contains continuous read coverage values for each bin, the Sei dataset contains binary labels for each bin corresponding to whether the bin overlapped a peak in each of the chromatin accessibility profiles. The Sei data were processed using 100-bp bins along the genome and using the hg38 reference genome.

### Single-cell kidney data

We obtained single-cell ATAC-sequencing data of primary human kidney tissue from three donors from [27]. These data had been clustered by [27] into 10 cell types, and pseudobulk ATAC data for each cell type had been generated. Peaks had also been grouped into disjoint clusters based on their accessibility profiles across cell types, giving the ubiquitous and cell type specific peak clusters used in our analyses. The data also included allele-specific chromatin accessibility information that we used in the allelic imbalance evaluations. All data were processed using the hg38 reference genome.

### Bulk immune cell data

We obtained bulk ATAC-sequencing data of 25 primary human immune cell types, sorted by flow cytometry, from four blood donors from [28]. ATAC-seq peaks had been grouped by [28] into disjoint clusters based on their accessibility profiles across cell types, giving the ubiquitous and cell type specific peak clusters used in our analyses. The data also included allele-specific chromatin accessibility information that we used in the allelic imbalance evaluations. All data were processed using the hg19 reference genome.

### Enformer predictions

The pretrained Enformer model was obtained from Avsec et al. [7]. To make predictions for a sequence, we averaged predictions over the forward and reverse complement sequence and minor sequence shifts to the left and right (1 nucleotide in each direction).

### Sei predictions

The pretrained Sei model was obtained from Chen et al. [8]. We used the *1 variant effect prediction.py* script in the Sei framework repository (https://github. com/FunctionLab/sei-framework) to make variant effect predictions for variants of interest.

### Partitioned heritability analyses

To assess trait heritability enrichment in cell type specific accessible regions within Enformer’s training data (684 accessibility tracks), we first obtained GWAS summary statistics for a set of UK Biobank traits that had previously been characterized to have heritability enrichment in the regions around tissue-specific genes [21]. After excluding any traits that did not pass quality control thresholds (i.e. low sample size, low confidence) on the Neale lab server, we retained a set of seven traits. Next, we grouped Enformer’s 684 accessibility tracks into 9 tissue categories mirroring the groupings in [21]. For each tissue category, we defined a set of “tissue peaks” as all peaks that were present in at least 30% of the corresponding accessibility tracks. We then divided these “tissue peaks” into high and low cell type specificity halves based on how many of the 684 accessibility tracks each peak was present in. We used stratified LD score regression (LDSC) [16] to measure trait heritability enrichment in the annotations corresponding to high and low cell type specificity peaks for each tissue category.

To assess whether variants with larger model-predicted differences in accessibility are enriched for trait heritability, we further subset the variants within the annotations described above (high and low cell type specificity peaks for each tissue category) based on Enformer’s variant effect predictions. Specifically, for each variant, we took the mean absolute SNP accessibility difference (SAD) score across all chromatin accessibility tracks corresponding to the tissue category. We then divided the variants within each annotation (e.g. high cell type specificity Cardiovascular peaks) into two equally sized groups based on the magnitude of their mean absolute SAD score to create two new annotations (i.e. “Low SAD score in high cell type specificity Cardiovascular peaks” and “High SAD score in high cell type specificity Cardiovascular peaks”). We again used stratified LDSC to measure trait heritability in high SAD score variants.

All partitioned heritability analyses were performed using stratified LDSC conditioned on all baselineLD v2.2 annotations that do not correspond to promoter and enhancer marks (baselineLD v2.2, https://alkesgroup.broadinstitute.org/ LDSCORE/) [39]. We do not condition on regulatory baseline annotations (i.e. promoter and enhancer marks) in our analyses to provide an unbiased estimate of heritability enrichment in ubiquitous versus cell type specific peaks, as the ubiquitous peaks in our datasets are likely to have more overlap with the regulatory baseline annotations.

### Fine-mapped GTEx eQTL classification

We obtained GTEx v8 eQTLs fine-mapped using the SuSiE method [40, 41] from the Supplementary Data in Avsec et al. [7]. Using these data, we evaluated Enformer and Sei on their ability to distinguish high posterior inclusion probability eQTLs (PIP *>* 0.9) from a matched negative set of low PIP eQTLs (PIP *<* 0.01). We used a similar methodology as in Avsec et al. [7] to perform the classification; in particular, we used model predictions from all chromatin accessibility tasks as features (684 features for Enformer; 2,372 features for Sei) and trained separate random forest classifiers for each tissue using eight-fold cross-validation. We used the default hyperparameters of scikit-learn and set the maximum features considered per decision tree split to *log*_2_ of the total number of features.

### Motif enrichment analysis

We assessed enrichment of TF motifs in cell type specific and ubiquitous peaks in the Loeb et al. [27] and Calderon et al. [28] datasets using SEA [42], for all TF binding profiles in the JASPAR 2022 CORE vertebrates non-redundant collection [33]. We used 50,000 randomly sampled 500bp genomic sequences as background, or control, sequences when looking for motif enrichment.

We defined putative CpG island peaks as peaks with greater than 50% GC content and a ratio greater than 0.6 of observed CpG dinucleotides versus the expected number based on the number of Gs and Cs in the peak. These criteria were based on the CpG islands UCSC genome browser track [43].

### CNN model architecture and training

We trained convolutional neural networks (CNNs) to map input DNA sequences (1344 bp) to continuous measures of chromatin accessibility (normalized ATAC-seq read counts). Our architecture is based on an updated version of the Basset model [2], which consists of 8 convolutional layers followed by two fully connected layers. For high capacity multi-task models, model capacity was increased by increasing the number of parameters in each layer. Specifications of the architecture of each model are provided in the supplementary data on Zenodo [32]. For all models, we modified the architecture to predict continuous – rather than binary – values, which has been shown to improve model generalizability and interpretability [11], and trained models to minimize the Poisson regression loss function. For all models, we used chromosomes 7, 14, and 15 for validation, chromosomes 4 and 5 for evaluation, and all other chromosomes for training. We used the Basenji repository [3] for data preprocessing, model training and evaluation.

### Evaluation of common training decisions

Using the model architecture and training scheme described above, we trained a suite of CNN models to evaluate the effect of common training decisions on cell type specific accessibility prediction. These include single-task versus multi-task learning, as well as increased capacity multi-task models. We also evaluated how training set composition impacts performance. A description of the 5 different training sets evaluated, which each have different compositions of peak versus non-peak sequences, is provided in Table 1. For each training decision, we trained three replicate models with different random initializations.

### Model-based transcription factor activity scores

Inspired by the motif insertion approach in Yuan and Kelley [44], we computed model-based transcription factor activity scores for the multi-task and single-task models for all TF binding profiles in the JASPAR 2022 CORE vertebrates non-redundant collection [33]. We first obtained 1,000 dinucleotide shuffled peak sequences from [44] as background sequences. For each TF and each background sequence, we sampled a motif sequence from the TF’s PWM and inserted it into the center of the background sequence. We made predictions for the background and motif-inserted sequences using each multi-task and single-task model, and took the difference in predicted accessibility between the motif-inserted and background sequences – averaged over the 1,000 sequences – as a model’s predicted TF activity score.

### Model-based TF motif discovery with TF-MoDISco

We used TF-MoDISco [29] to identify TF motifs driving predictions in cell type specific and ubiquitous peaks for the multi-task and single-task models. For a sampled set of sequences in the ubiquitous and cell type specific peak clusters, we computed model attribution scores for each model variant (i.e. baseline multi-task, high capacity multi-task, single-task) and cell type using gradient*input. We then ran TF-MoDISco using the tfmodisco-lite implementation (https://github.com/jmschrei/tfmodisco-lite) to identify seqlets with high attribution scores and their similarity to known TF motifs in the JASPAR 2022 CORE vertebrates non-redundant collection [33]. We considered all motif matches with a q-value *<* 0.05 when computing the Jaccard similarity in motif matches between cell types.

---

**Fig. S1.**
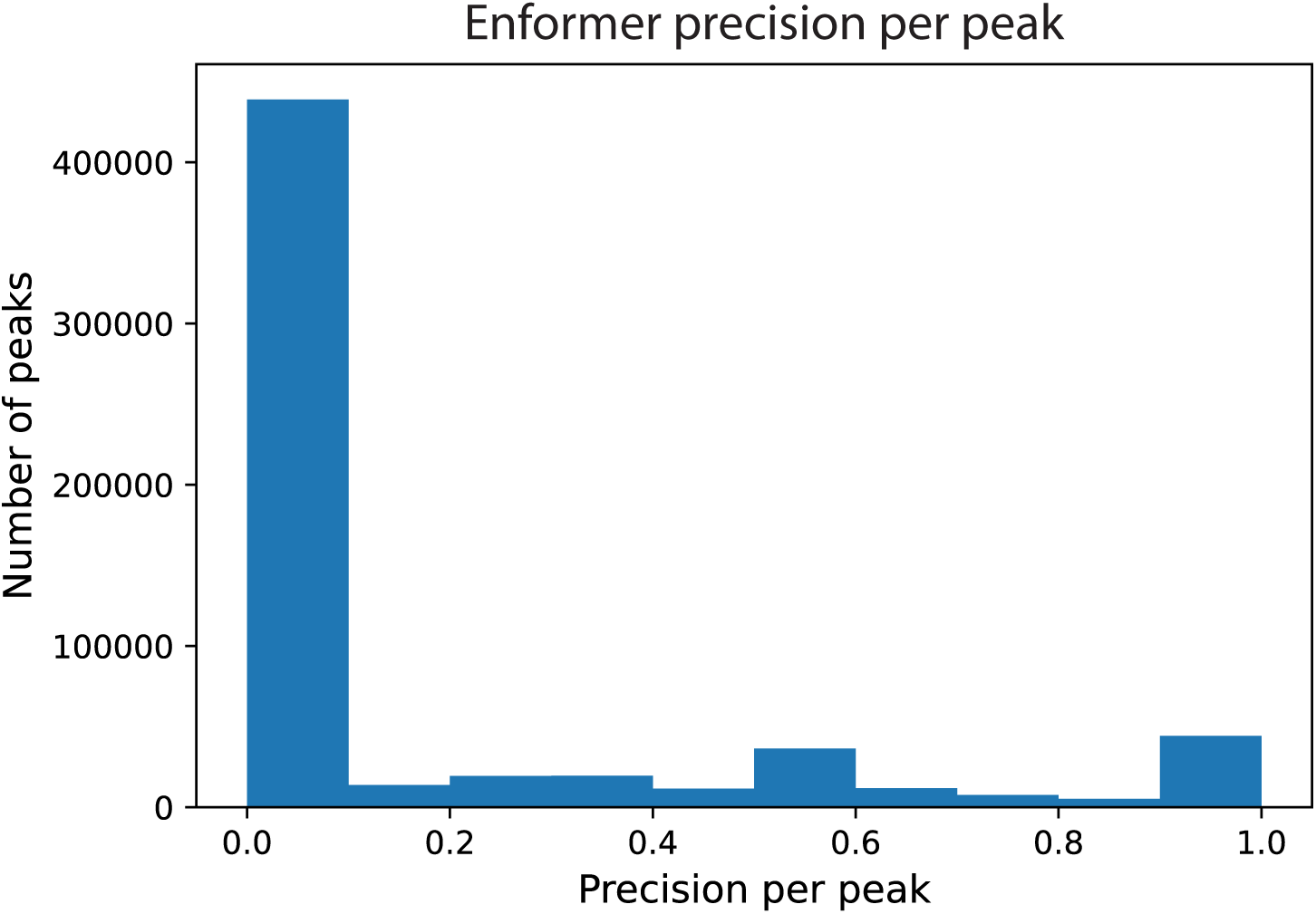
Cell type precision of Enformer’s chromatin accessibility predictions. For all sequences in Enformer’s test set that are a peak in at least one experimental chromatin accessibility track, we compute the precision of Enformer’s predictions across chromatin accessibility tracks. For each peak sequence, we first identify the true number (*N*) of experimental chromatin accessibility tracks with a peak. Then, based on the *N* chromatin accessibility tracks Enformer predicts to have the highest accessibility for that peak, we compute the precision of its prediction. To ensure an even comparison across tracks, we compare z-score normalized predictions for each track.

**Fig. S2.**
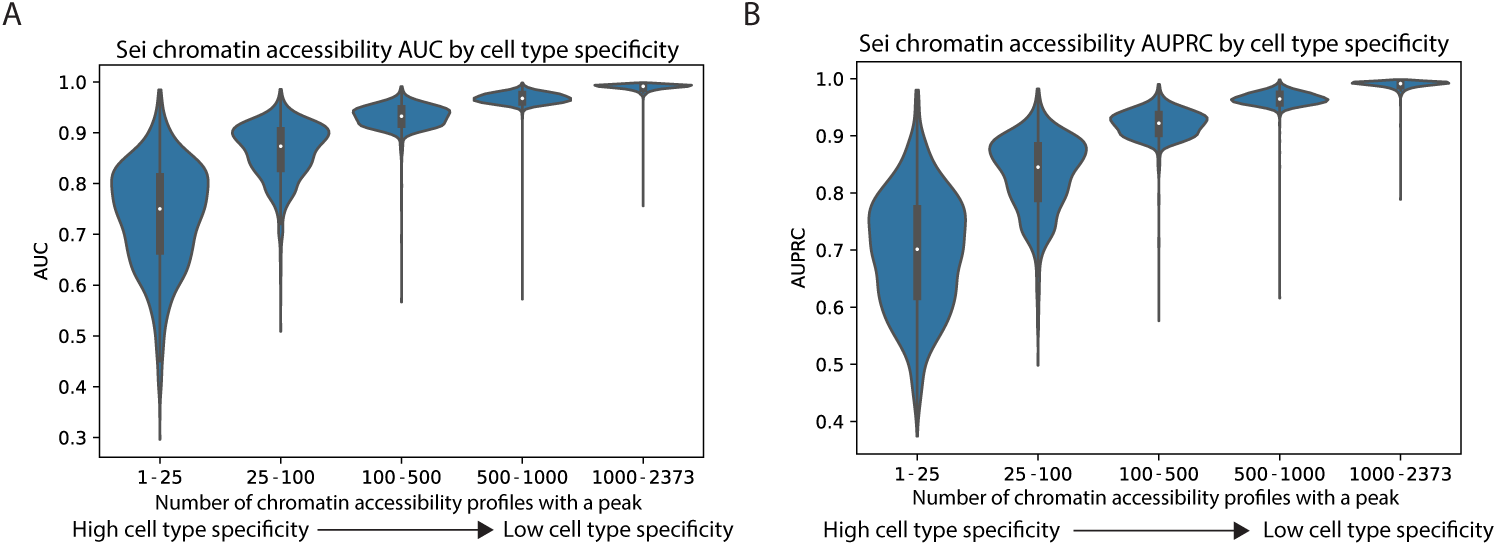
Sei chromatin accessibility prediction performance stratified by cell type specificity. Sei’s chromatin accessibility prediction A) AUROC and B) AUPRC measured for sequences in accessibility peak bins with varying degrees of cell type specificity. Sei’s chromatin accessibility predictions are highly accurate in low cell type specificity peaks and less accurate in high cell type specificity peaks.

**Fig. S3.**
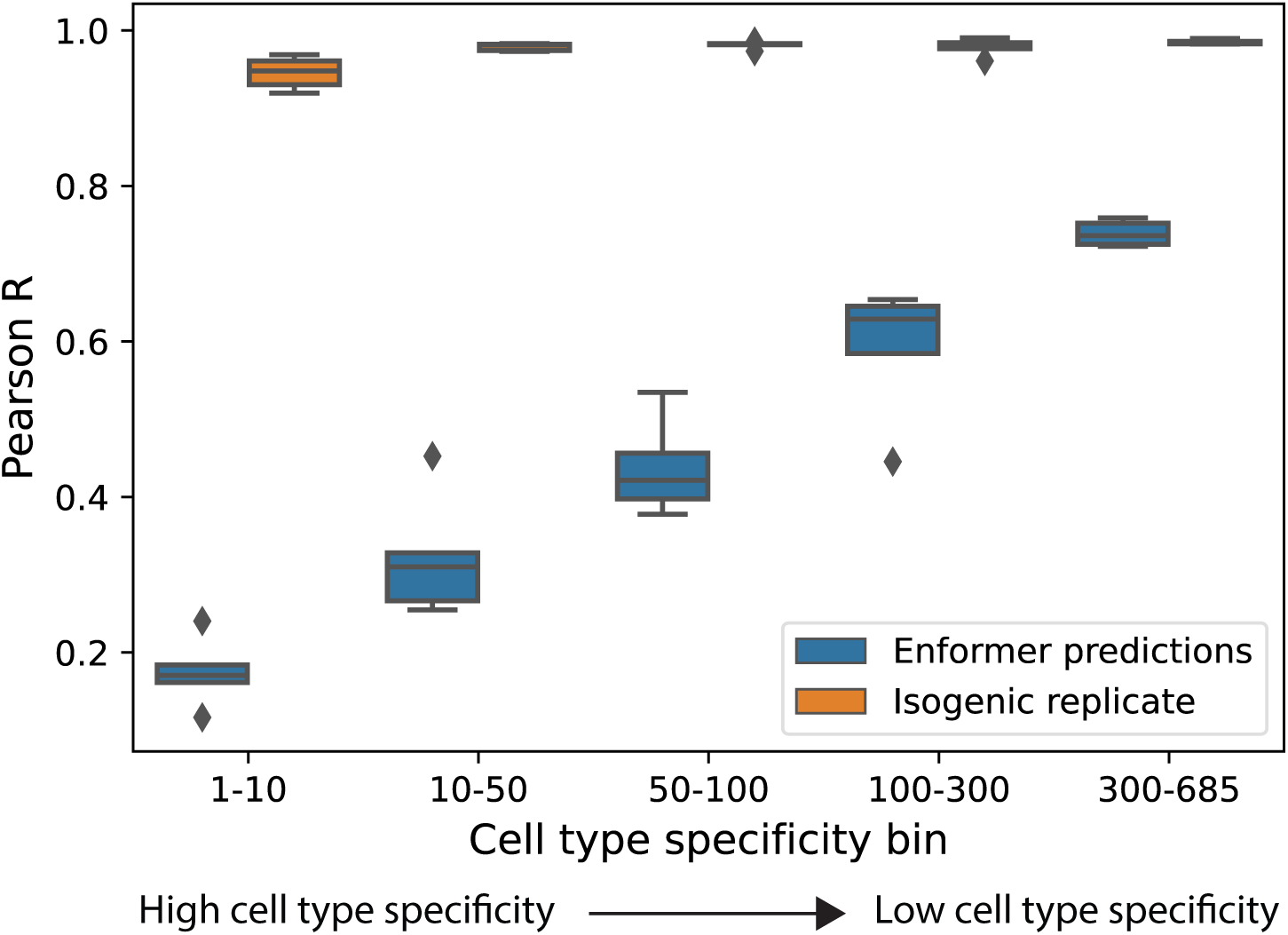
Comparison of Enformer prediction performance versus isogenic replicate correlation for peaks with varying degrees of cell type specificity. For five representative Enformer DNase tracks with isogenic replicate data available on ENCODE, the peak height correlation between experimental replicates remains high even for high cell type specificity peaks. Thus experimental noise does not explain the dramatic drop in Enformer’s predictive performance for cell type specific peaks.

**Fig. S4.**
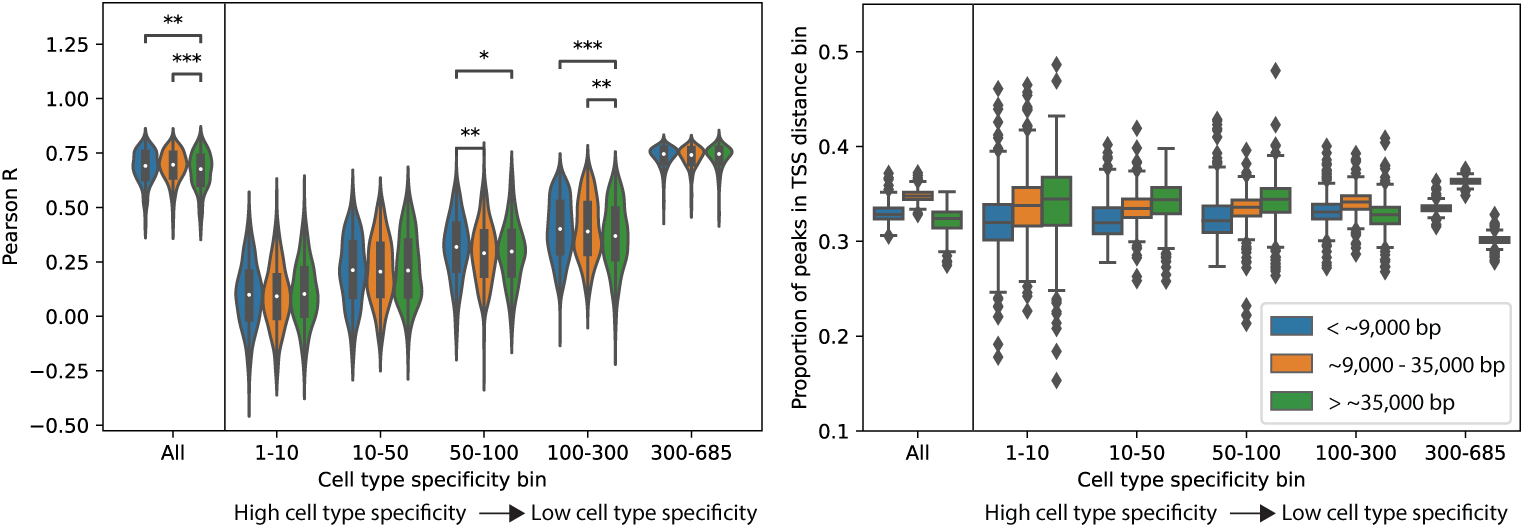
Enformer chromatin accessibility prediction performance stratified by cell type specificity and TSS distance. A) Enformer prediction performance for peaks stratified by TSS distance only (left, ”All”) and for peaks stratified by both TSS distance and cell type specificity. B) The relative proportion of proximal versus distal sequences in each cell type specificity bin.

**Fig. S5.**
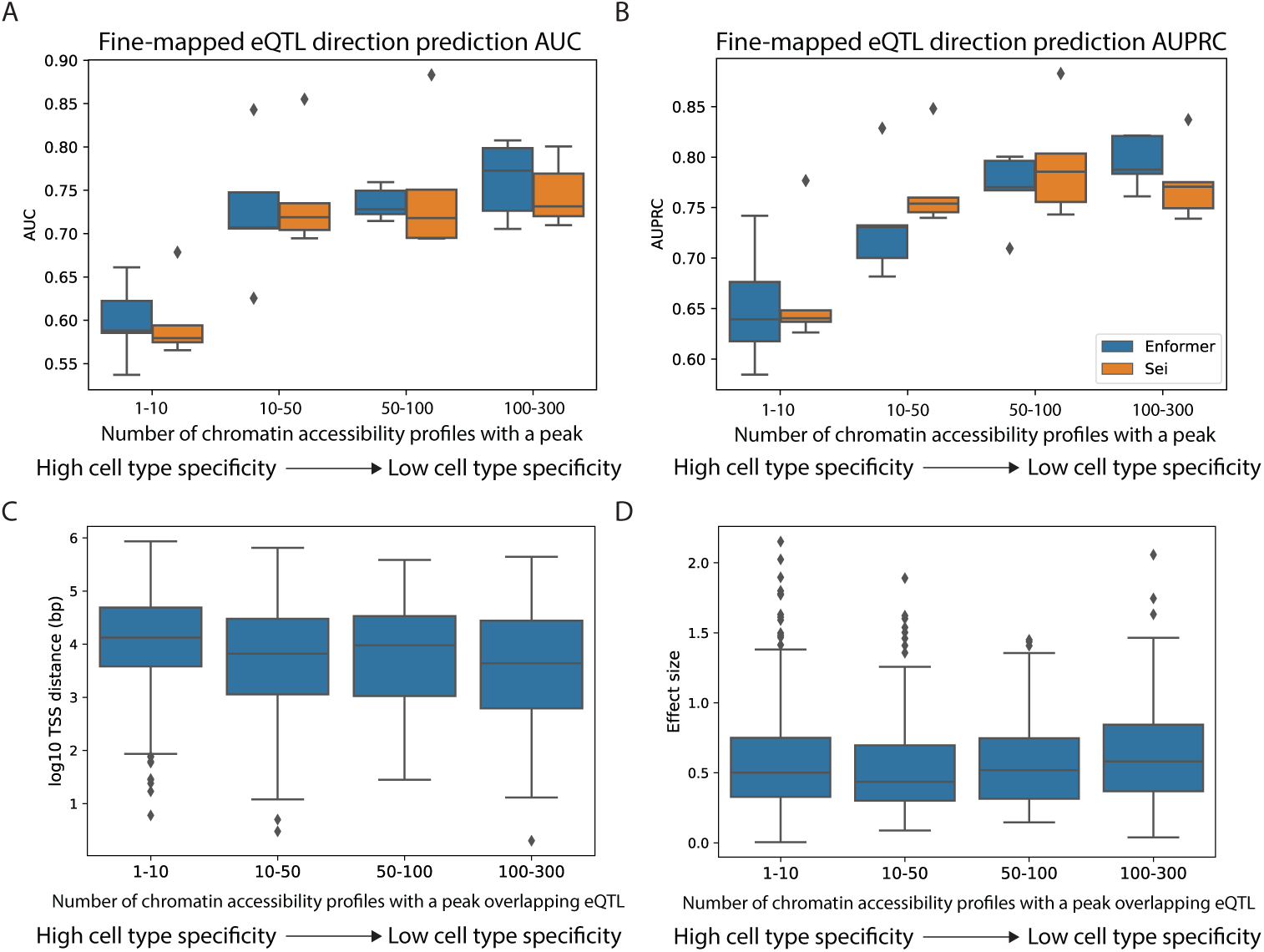
Enformer and Sei eQTL direction prediction performance, TSS distance distribution, and effect size distribution stratified by cell type specificity. Enformer and Sei’s eQTL direction prediction A) AUROC and B) AUPRC measured for high posterior inclusion probability eQTLs (PIP *>* 0.9) residing in accessibility peaks with varying degrees of cell type specificity. C) Distance to the nearest gene transcription start site (TSS) distributions and D) eQTL effect size distributions for high PIP eQTLs in each cell type specificity bin.

**Fig. S6.**
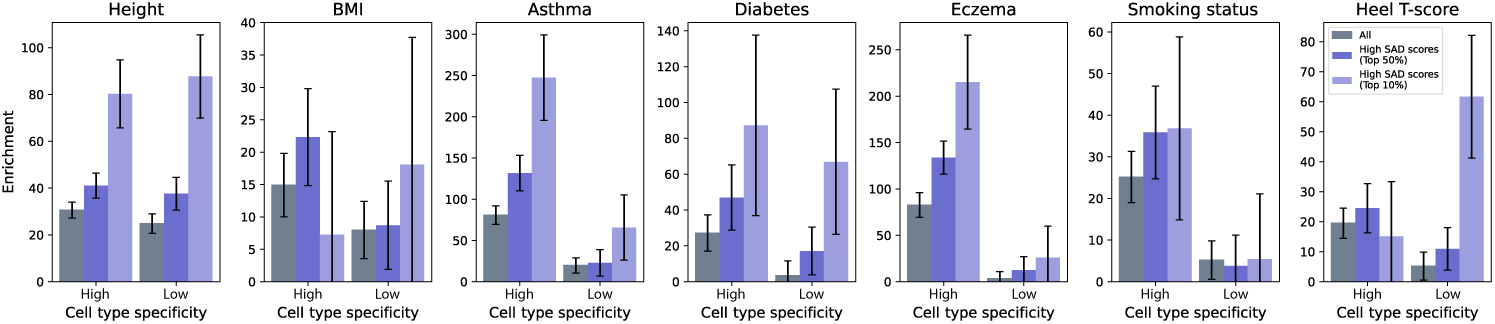
Trait heritability enrichment of Enformer high SAD score variants is robust to choice of threshold. To verify that the results in Fig. 2D are robust to the choice of threshold, here we subset variants by taking only the top 10% highest Enformer SAD scores and assess enrichment of trait heritability in this subset.

**Fig. S7.**
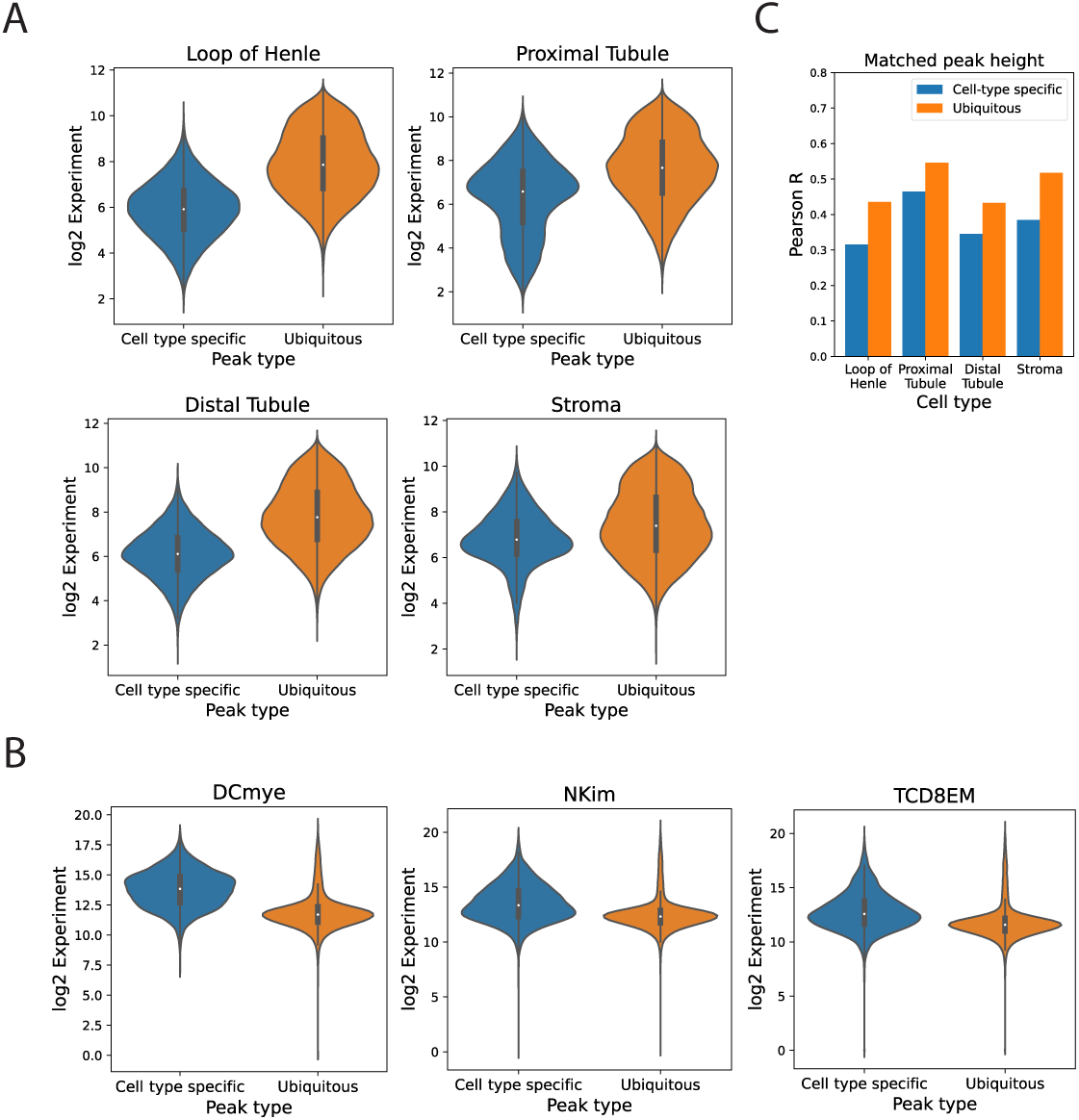
Chromatin accessibility prediction performance in peak-height matched cell type specific and ubiquitous peaks. Peak height distributions for cell type specific and ubiquitous peaks in the A) Loeb et al. [27] and B) Calderon et al. [28] data. C) Predictive performance (reference accuracy) in cell type specific and ubiquitous peaks in the Loeb et al. [27] data after matching on peak height. Reference accuracy is measured as the Pearson correlation between experimentally measured versus predicted accessibility.

**Fig. S8.**
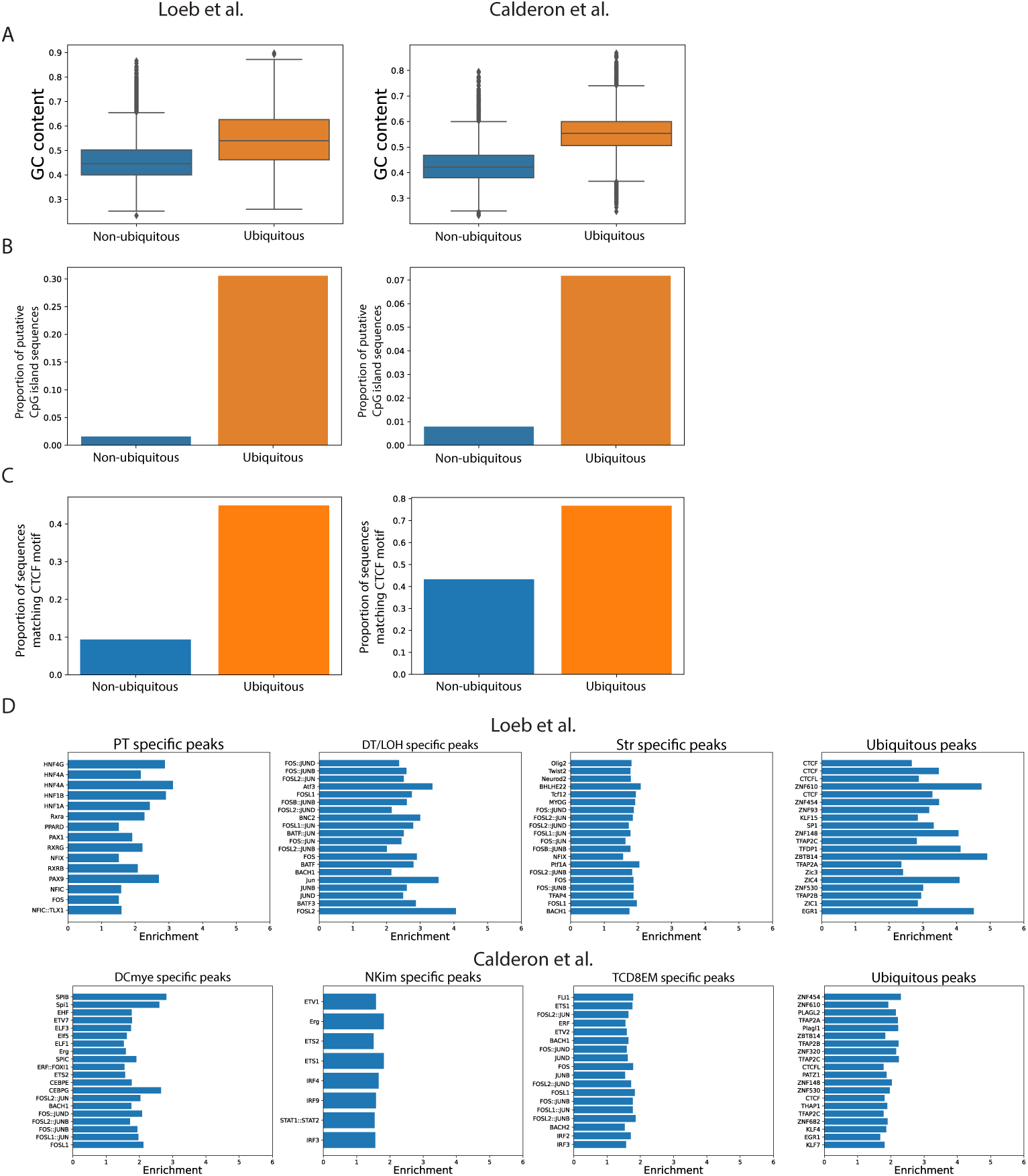
Sequence features and top enriched motifs of ubiquitous and non-ubiquitous peaks in the Loeb et al. [27] and Calderon et al. [28] data. A) GC content fraction for sequences in ubiquitous and non-ubiquitous peaks. B) Proportion of sequences containing putative CpG islands, and C) proportion of sequences containing CTCF motifs in ubiquitous and non-ubiquitous peaks. D) Top motifs enriched in cell type specific and ubiquitous peaks relative to non-peaks. We show the enrichment of the top motifs that are present in at least 5% of the peaks in each peak set, have an enrichment p-value *≤* 10*^−^*^6^, and an enrichment greater than 1.5.

**Fig. S9.**
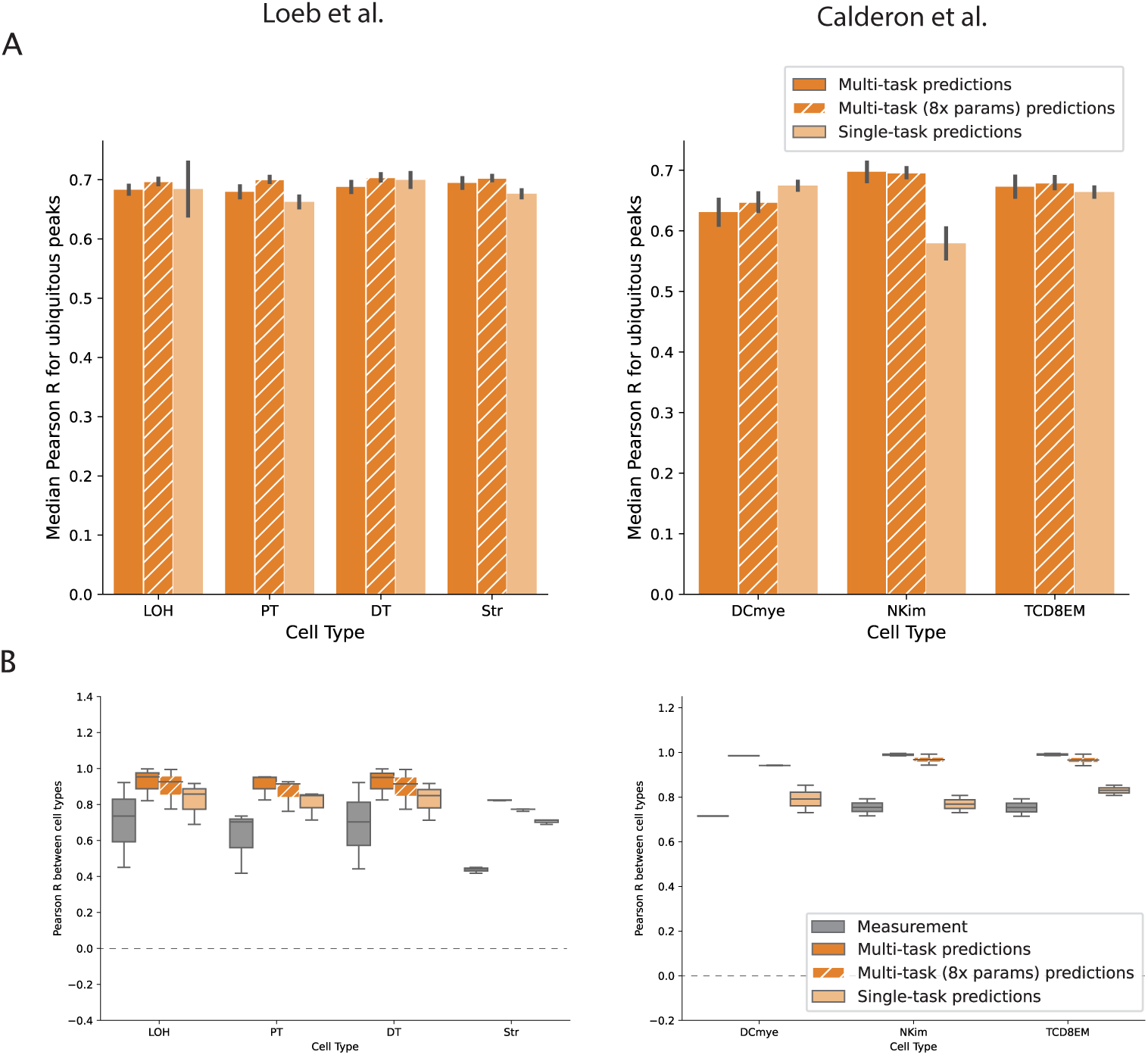
Comparison of multi-task versus single-task model performance in ubiquitous peak regions. A) Reference accuracy of multi-task versus single-task models evaluated in ubiquitous peak regions. B) Pairwise correlations of peak height between cell types for experimental (gray) and model predicted accessibility (dark and light orange) in ubiquitous peaks. Model predicted accessibility is more correlated across cell types than experimentally measured accessibility, and this overcorrelation is slightly more pronounced in predictions from multi-task models than predictions from single-task models.

**Fig. S10.**
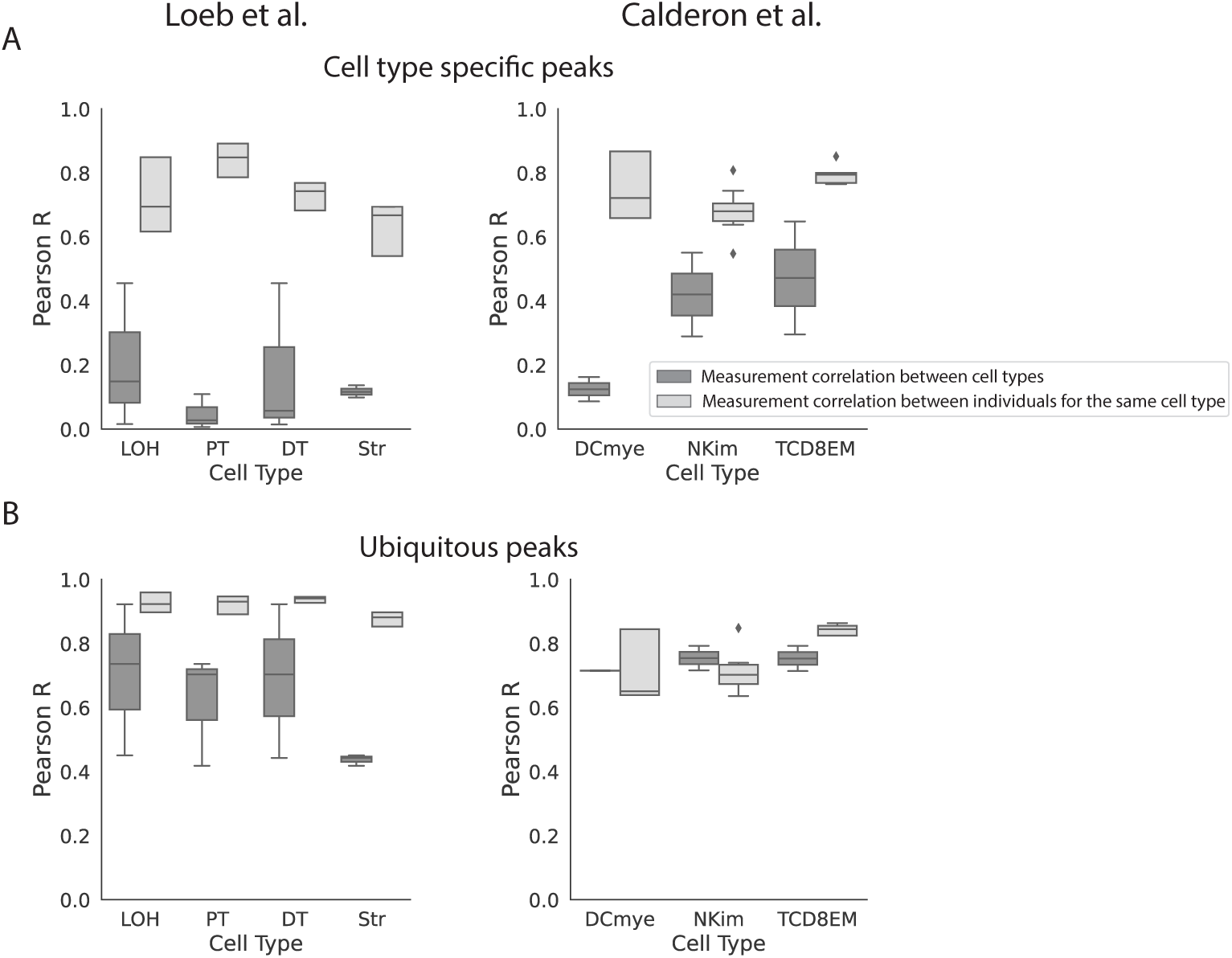
Experimental measurement noise in cell type specific peaks does not explain the over-correlation in model predictions between cell types. A) Cell type specific peaks have low peak height correlation across cell types (dark gray), but high correlation across biological replicates (light gray). We note that because these biological replicates represent samples from different donors, they encompass both biological and experimental sources of variability and represent an upper bound on experimental noise. Thus experimental noise in cell type specific peaks does not explain the observed over-correlation of the model predictions across cell types (Fig. 4B). B) Peak heights of ubiquitous peaks are highly correlated both across cell types (dark gray) and across biological replicates (light gray).

**Fig. S11.**
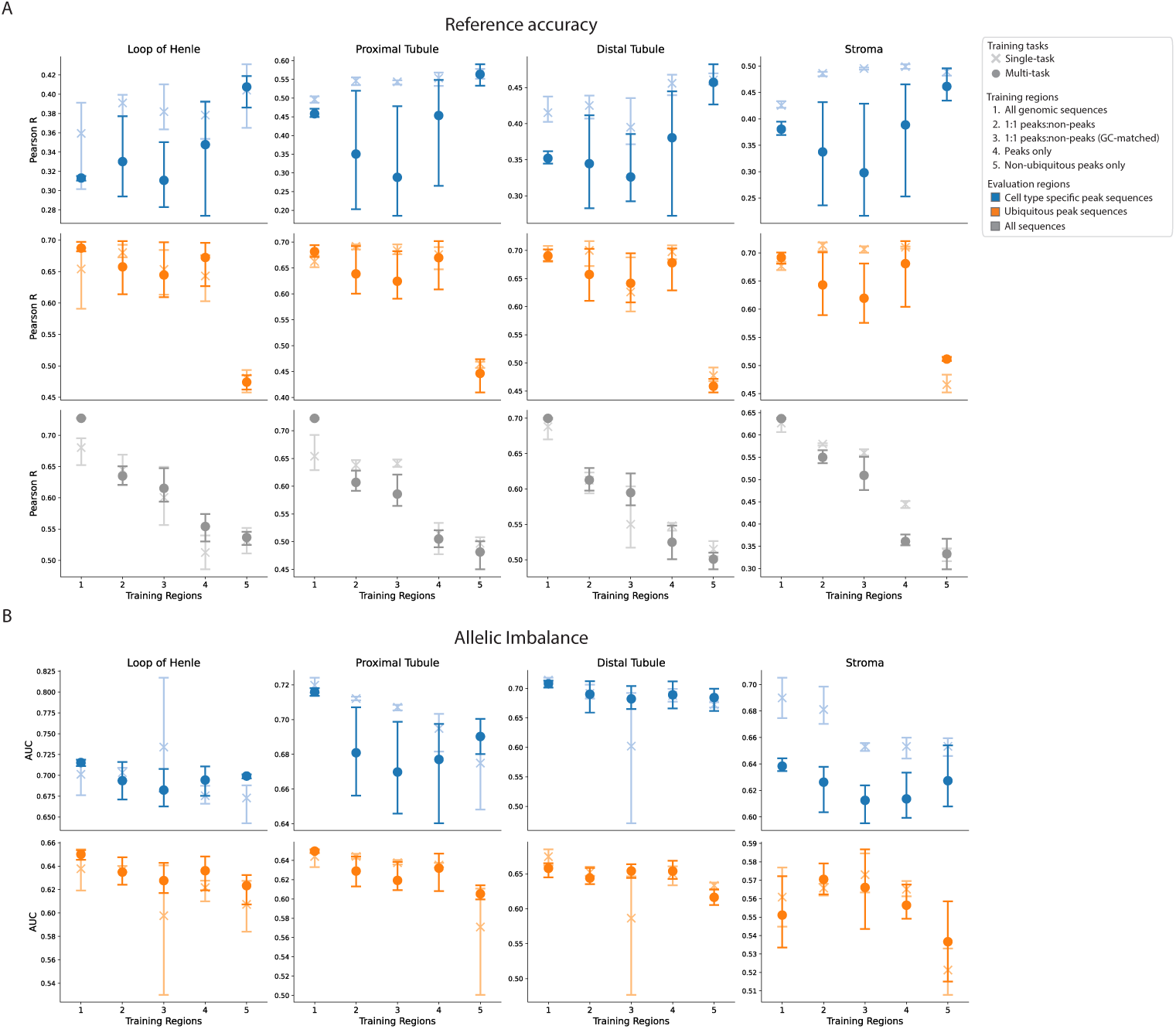
Evaluating single-task versus multi-task learning and training set composition in the Loeb et al. [27] data. A) Reference accuracy (Pearson correlation between experimentally measured and predicted accessibility) and B) variant effect accuracy (AUROC for the model’s ability to predict the direction of imbalance, or the higher accessibility allele, based on allelic imbalance measurements) in ubiquitous and cell type specific peak regions for the different modeling decisions that we evaluated.

**Fig. S12.**
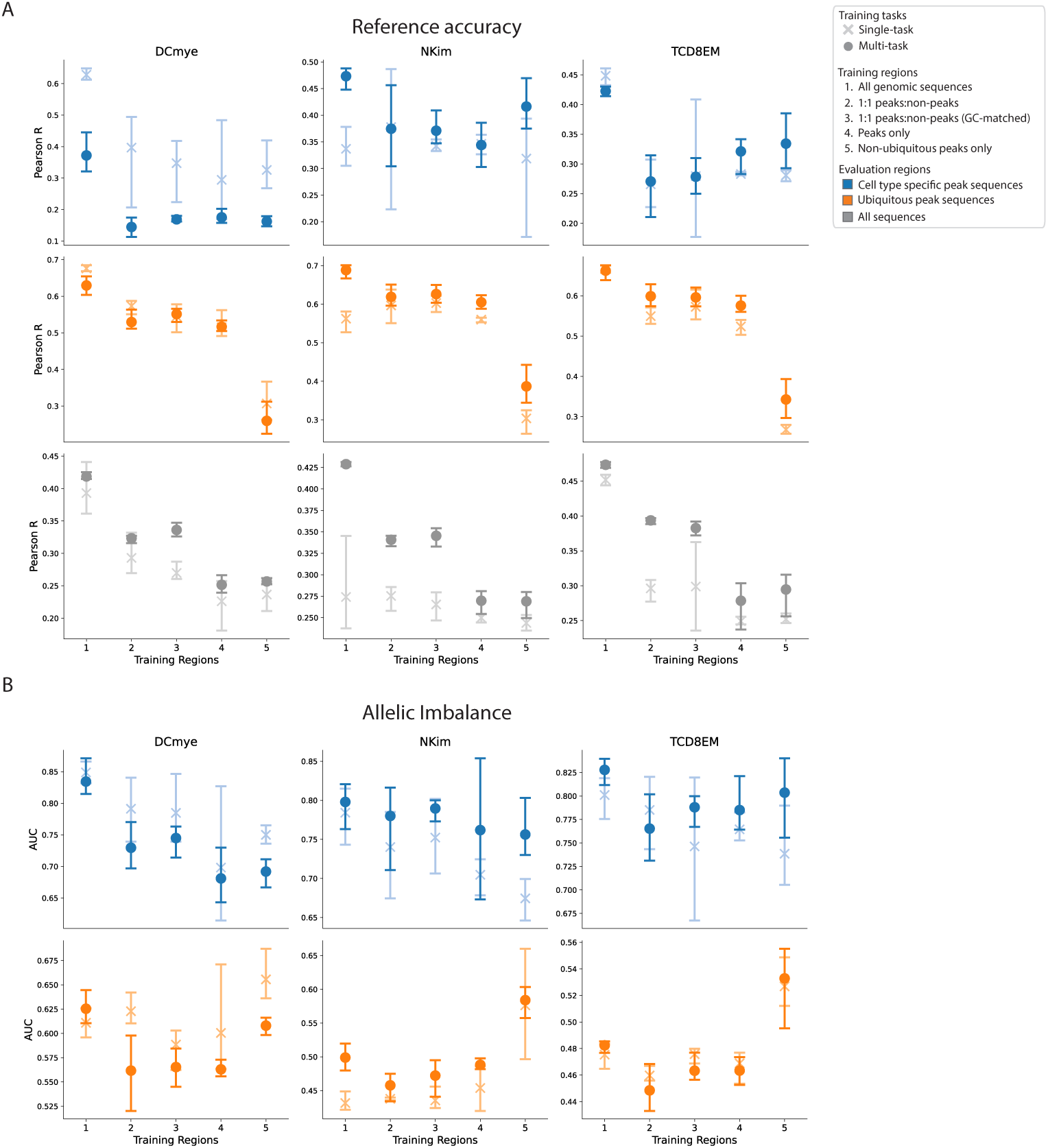
Evaluating single-task versus multi-task learning and training set composition in the Calderon et al. [28] data. A) Reference accuracy (Pearson correlation between experimentally measured and predicted accessibility) and B) variant effect accuracy (AUROC for the model’s ability to predict the direction of imbalance, or the higher accessibility allele, based on allelic imbalance measurements) in ubiquitous and cell type specific peak regions for the different modeling decisions that we evaluated.

**Fig. S13.**
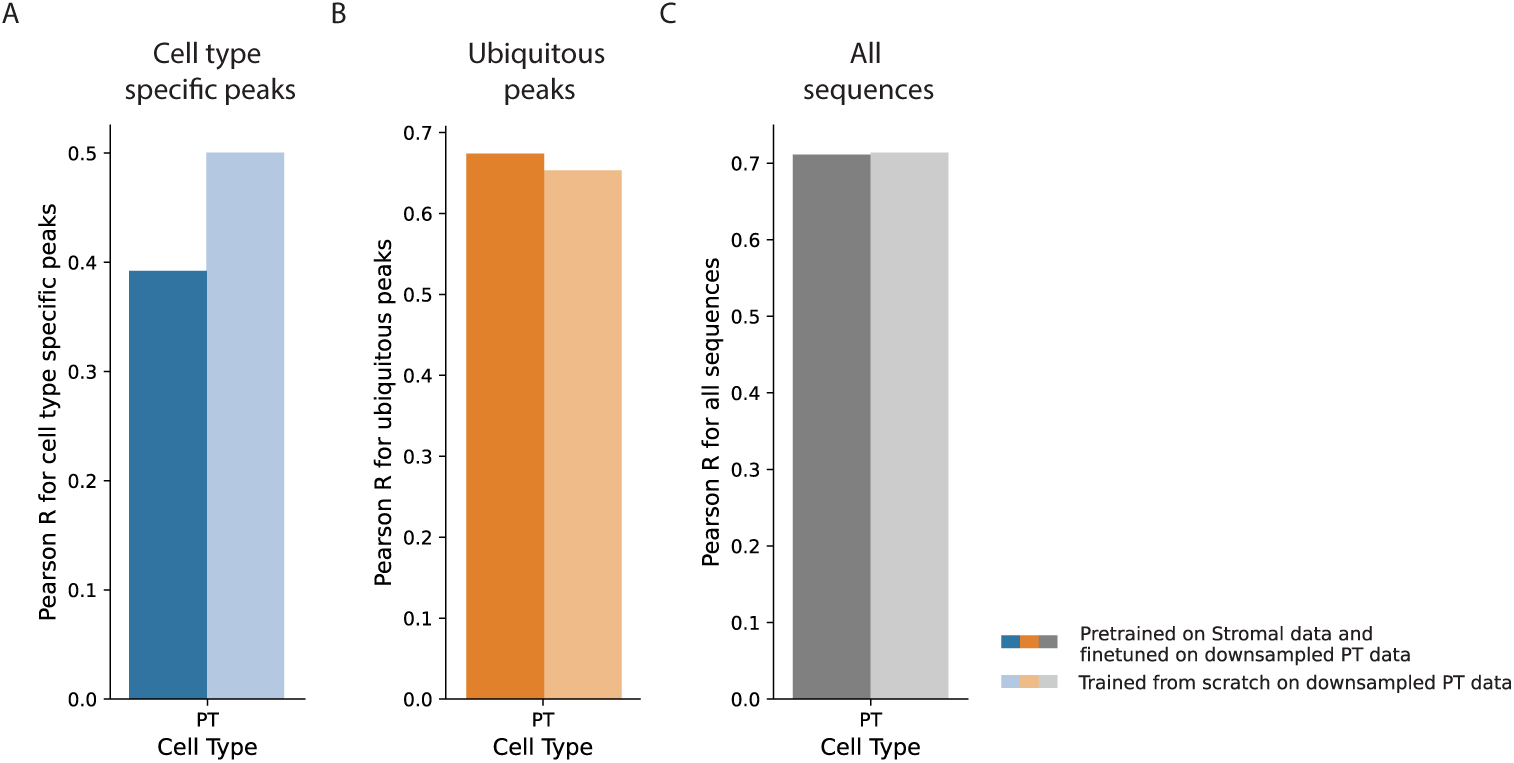
Evaluation of a transfer learning strategy on cell type specific and ubiquitous peak prediction. Using the Loeb et al. [27] data, we evaluate a transfer learning strategy where a single-task model is first trained on a cell type with abundant high-quality data, and then fine-tuned on a cell type with lower quality data, reasoning that for cell types with lower quality data, this approach may retain beneficial aspects of both multi-task and single-task models. Specifically, we trained two additional model variants using a downsampled version of the Proximal Tubule data (downsampled to approximately 10% of the original number of collected cells): 1) a single-task model trained from scratch on the downsampled Proximal Tubule data (lighter colors), and 2) a single-task model pretrained on the Stromal cell data, and fine-tuned on the downsampled Proximal Tubule data (darker colors). We show the performance of both models on A) cell type specific peaks, B) ubiquitous peaks, and C) all genomic sequences.

**Fig. S14.**
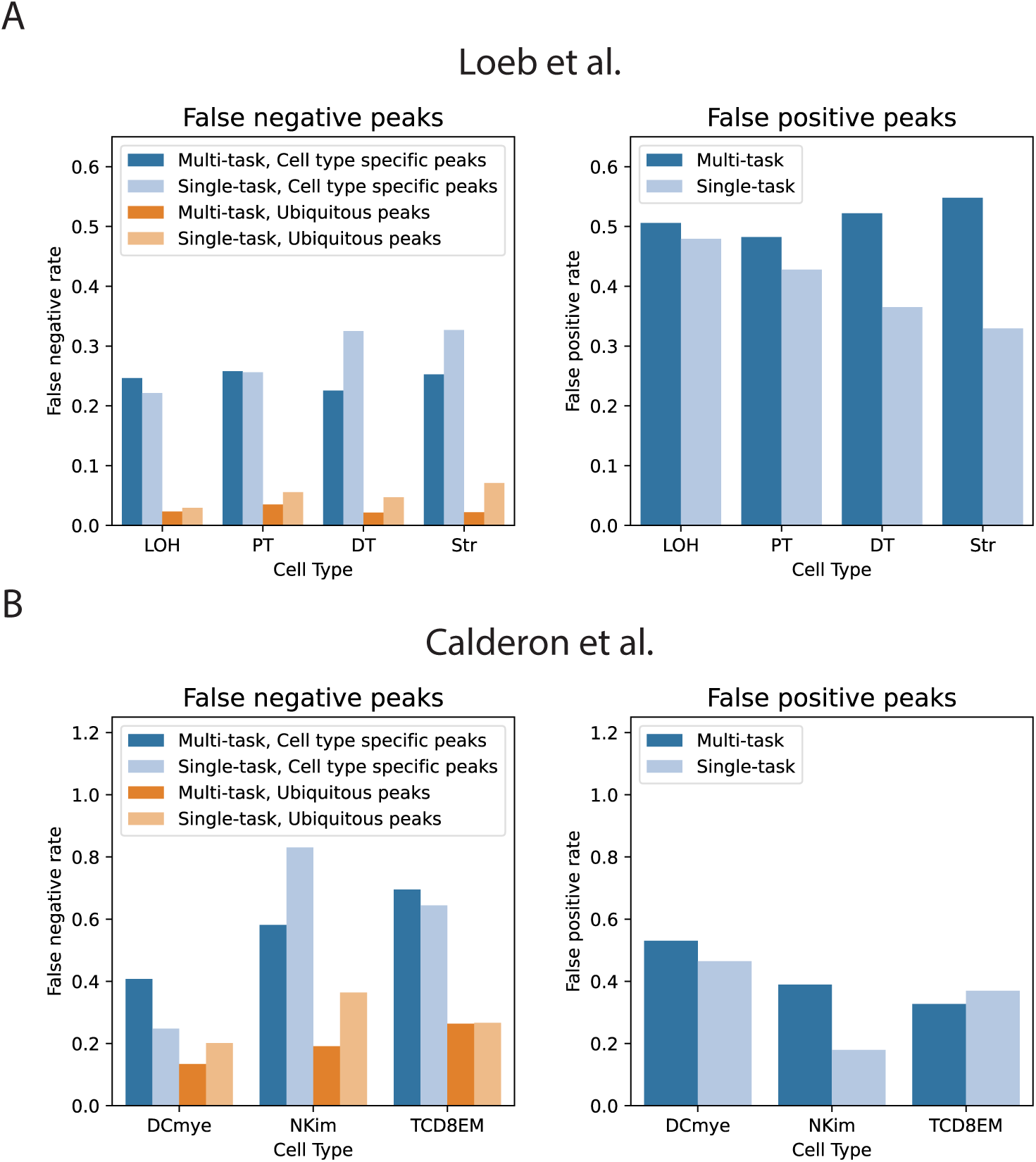
Comparison of multi-task and single-task model false negative and false positive peak prediction rates. For both the A) Loeb et al. [27] and B) Calderon et al. [28] datasets, we compare the false negative and false positive rate of predictions from the baseline multi-task and single-task models. The baseline multi-task models tend to have higher false positive rates than single-task models, meaning that they predict that a peak is accessible in cell types in which the peak is not accessible. For both types of models, the false negative rate is higher for cell type specific peaks than ubiquitous peaks.

**Fig. S15.**
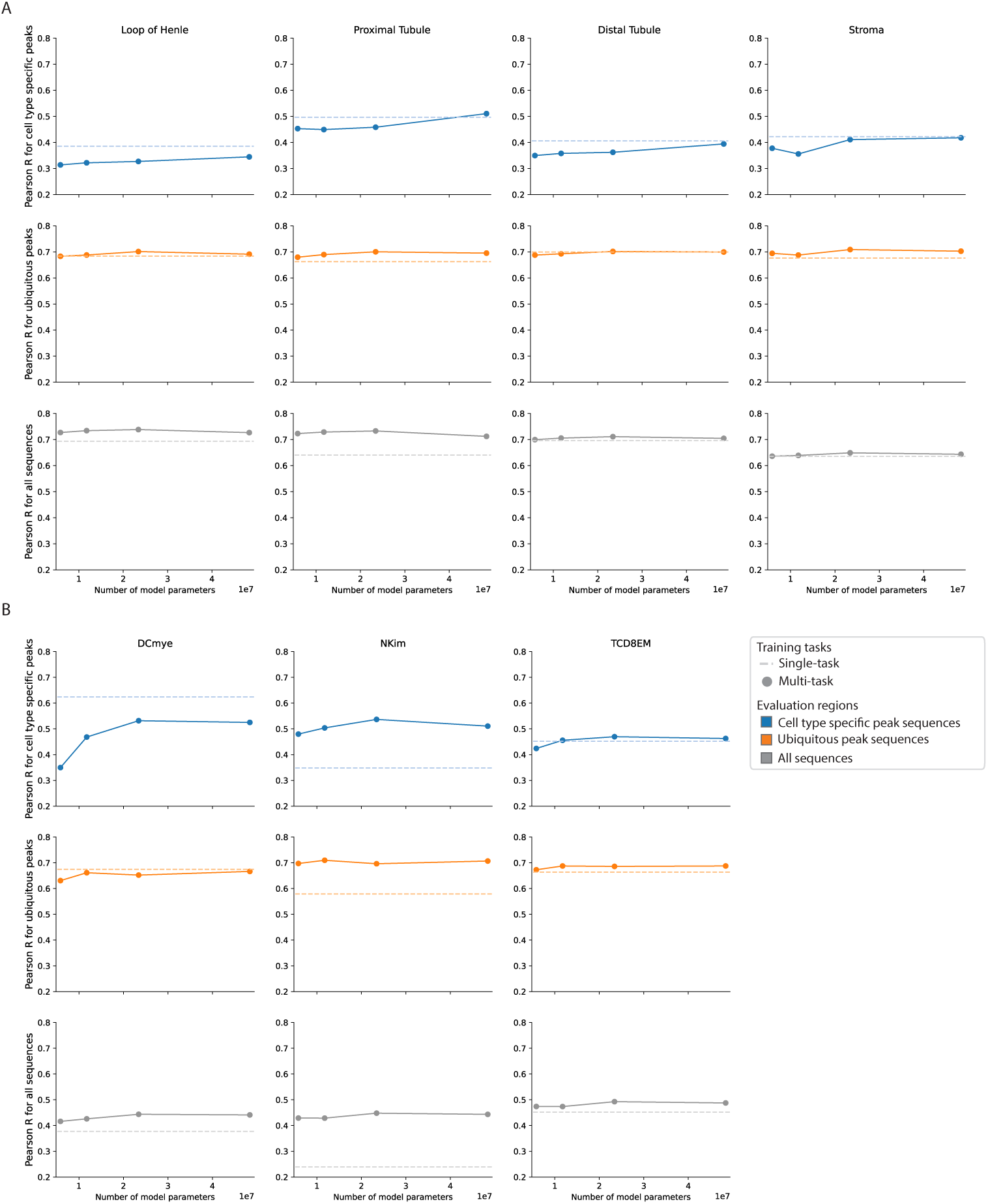
Increased multi-task model capacity improves cell type specific accessibility prediction. Analysis of the effect of increasing multi-task model capacity on prediction performance in cell type specific peaks, ubiquitous peaks, and all genomic sequences using the A) Loeb et al. [27] and B) Calderon et al. [28] data. For each cell type, performance of a single-task model with a similar number of parameters as the baseline (smallest) multi-task model is shown with a dashed line. Model capacity for the multi-task models was increased by increasing the number of parameters in each layer.

**Fig. S16.**
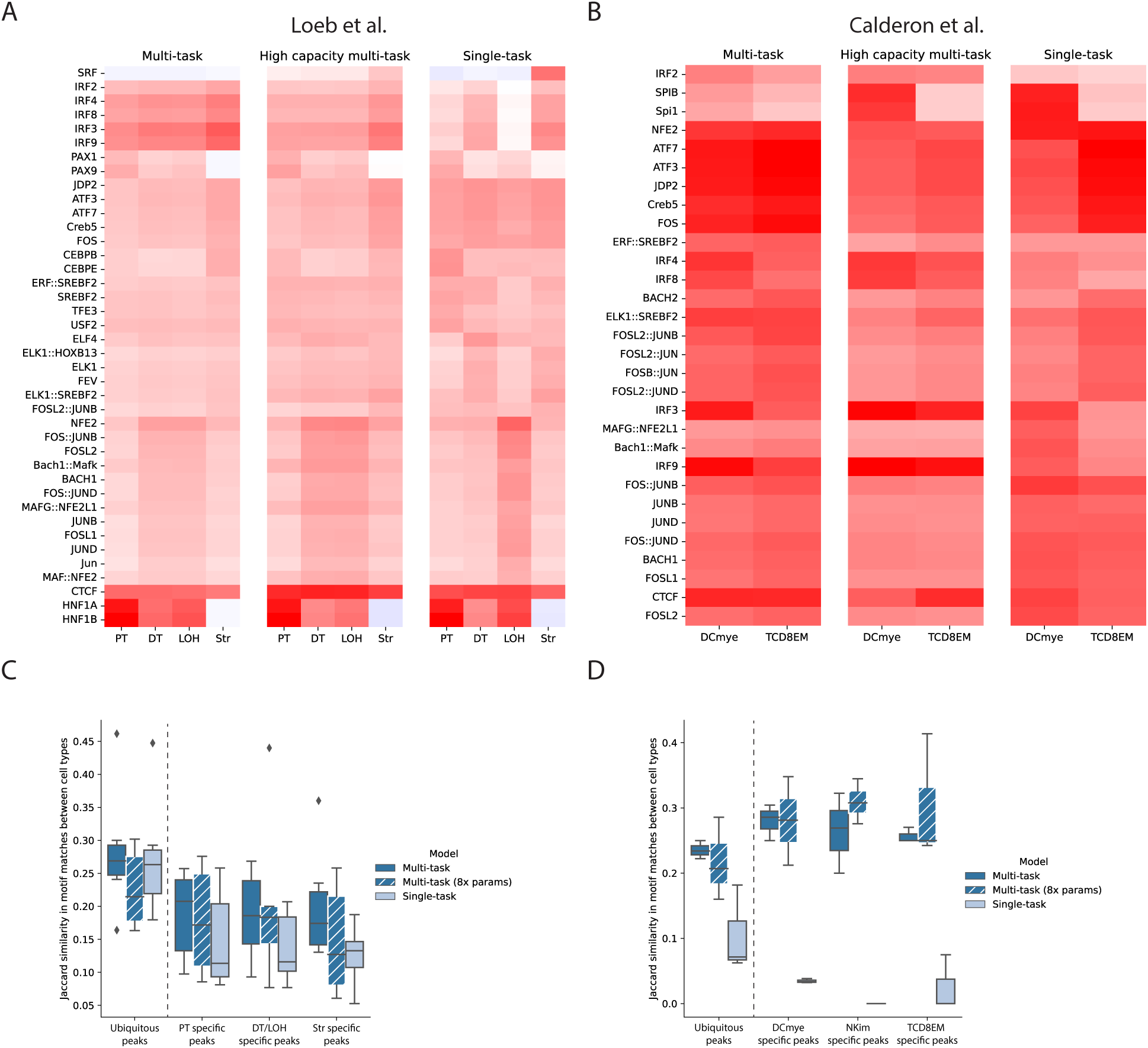
Interpretability analysis of TF motifs learned by single-task versus multi-task models. We compute model predicted TF activity scores for both the A) Loeb et al. [27] and B) Calderon et al. [28] datasets in each cell type for baseline multi-task, high capacity multi-task, and single-task models. TFs with the greatest model predicted activity (Z-score *≥* 3) are plotted. TF activity scores were computed by comparing model predicted activity for a set of dinucleotide shuffled background sequences versus the dinucleotide shuffled background sequences with the TF’s canonical motif inserted at the center of the sequence. For both the C) Loeb et al. [27] and D) Calderon et al. [28] datasets, we also use TF-MoDISco to identify the TF motifs driving predictions in ubiquitous and cell type specific peaks. For the baseline multi-task, high capacity multi-task, and single-task models, we report the Jaccard similarity in the motifs identified by TF-MoDISco across cell types.

**Fig. S17.**
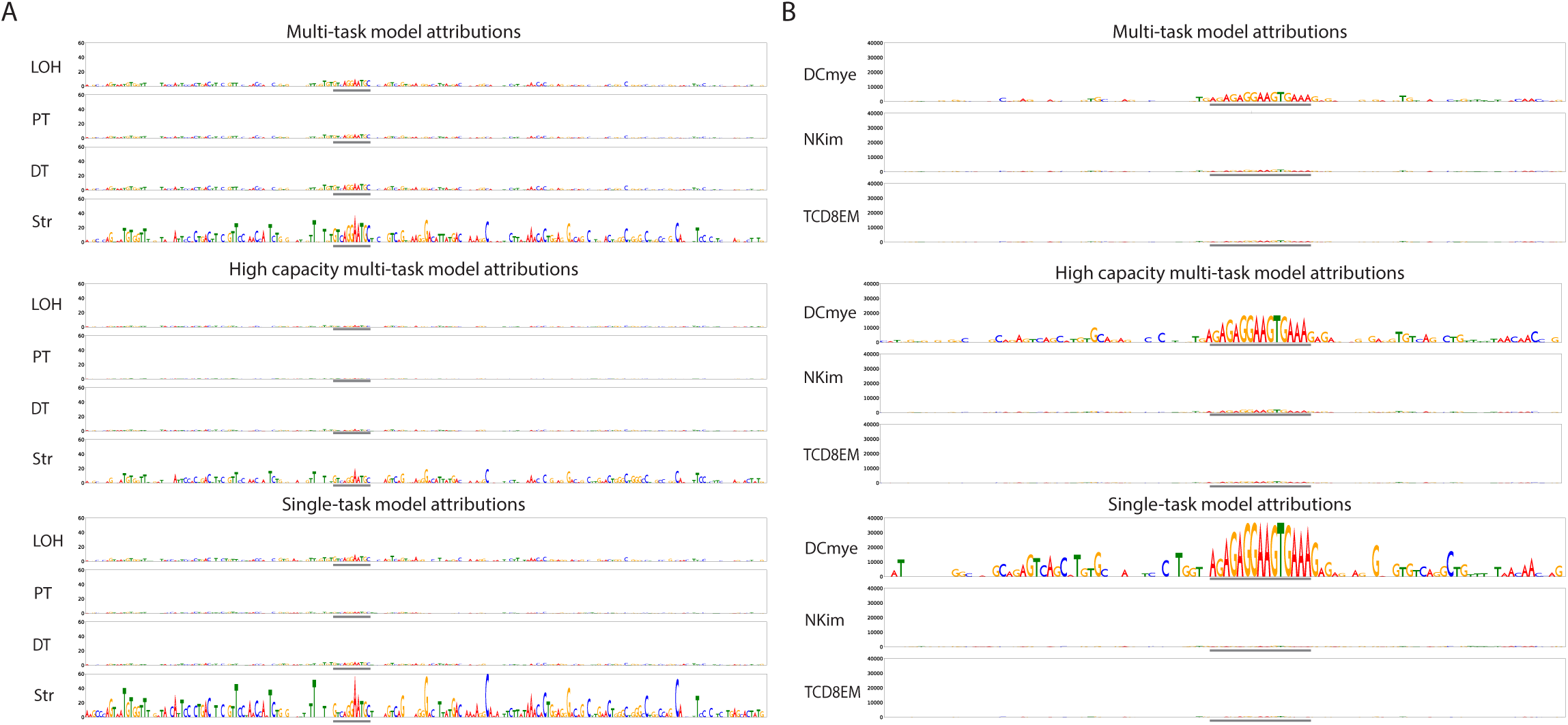
*In silico* mutagenesis of cell type specific peaks for tissue-specific models. A) *In silico* mutagenesis (ISM) of a Stromal cell specific peak – chr5:68,137,529-68,137,729 (hg38 coordinates) – that is mispredicted by the baseline multi-task model to be a peak in additional cell types, but is correctly predicted by the single-task models to only be accessible in Stromal cells. The ISM score for each position is the maximum absolute decrease in predicted accessibility over all non-reference nucleotides compared to the reference nucleotide. ISM reveals that a TEAD-like motif is contributing to the predictions of the baseline multi-task model in both Stromal and non-Stromal cell types, while this TEAD-like motif is less apparent in non-Stromal cell types for the high-capacity multi-task and single-task models. The location of the TEAD-like motif is indicated with gray bars underneath the ISM tracks. ISM scores are plotted on the same scale for all models. B) ISM of a dendritic cell specific peak – chr5:73,748,769-73,748,869 (hg19 coordinates) – that is mispredicted by the baseline multi-task model to be a peak in additional cell types, but is correctly predicted by the single-task models to only be accessible in dendritic cells. ISM reveals a SPIB-like motif weakly informing predictions of the baseline multi-task model, and more strongly driving dendritic cell predictions for the high-capacity multi-task and single-task models. The location of the SPIB-like motif is indicated with gray bars underneath the ISM tracks. ISM scores are plotted on the same scale for all models.

**Fig. S18.**
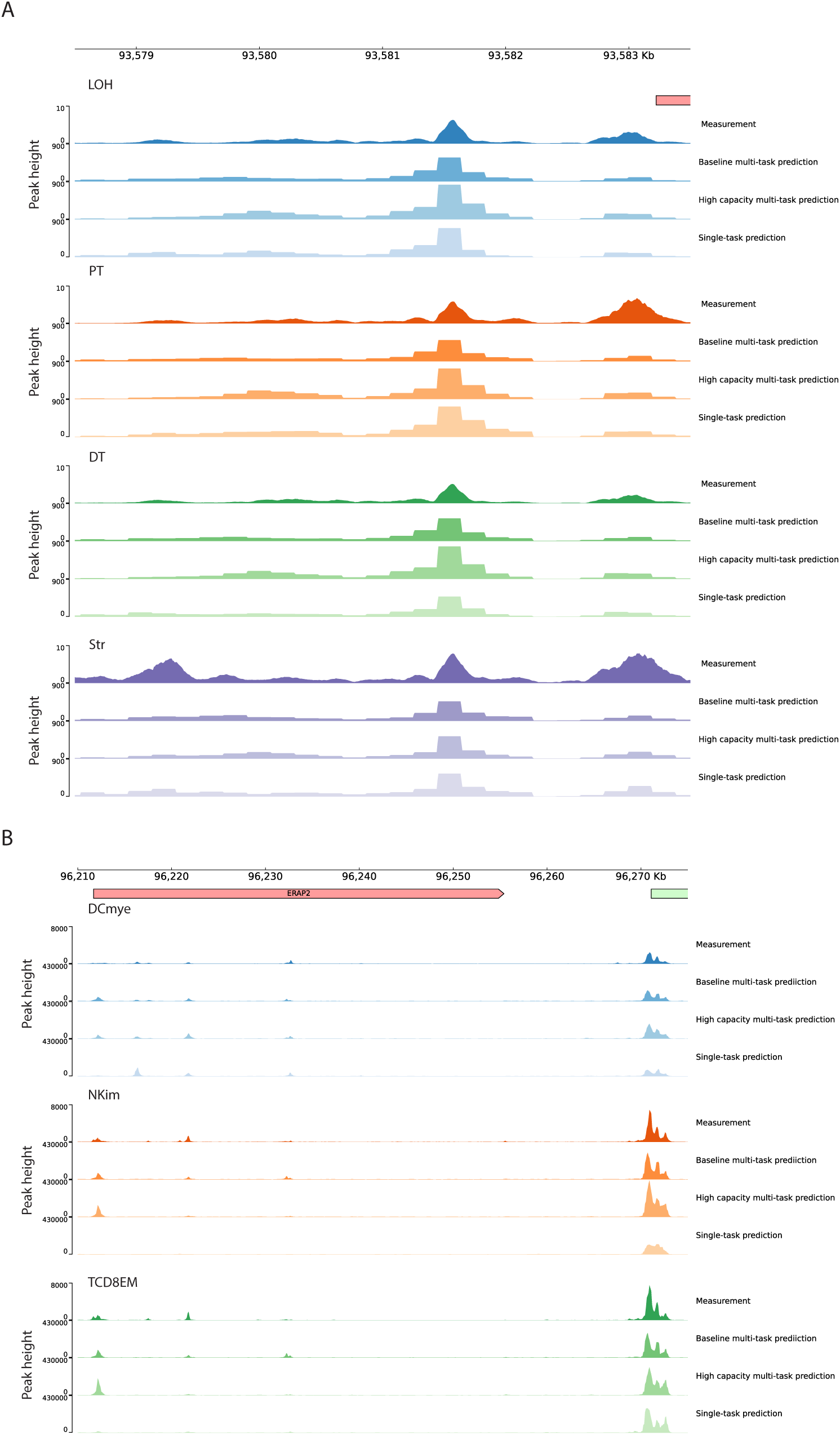
Single-task versus multi-task model predictions at the *NR2F1* and *ERAP2* loci. A) Experimentally measured accessibility in the Loeb et al. [27] data and predicted accessibility profiles from baseline multi-task, high capacity multi-task, and single-task models for the region around *NR2F1*. B) Experimentally measured accessibility in the Calderon et al. [28] data and predicted accessibility profiles from baseline multi-task, high capacity multi-task, and single-task models for the region around *ERAP2*.

**Fig. S19.**
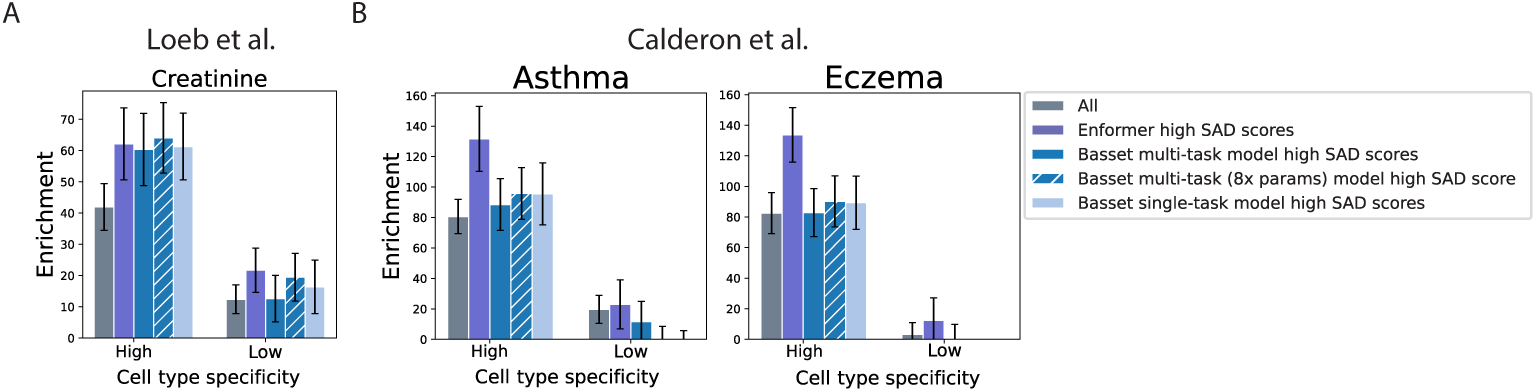
Comparing variant effect predictions from tissue-specific models and Enformer on GWAS heritability enrichment. For single-task and multi-task models trained on the A) Loeb et al. [27] and B) Calderon et al. [28] data, as well as Enformer, we subset variants in high and low cell type specificity peak regions based on each model’s SNP Accessibility Difference (SAD) scores and assess enrichment of trait heritability for tissue-matched traits. We use the median SAD score for all variants in a particular peak set (e.g.“Kidney high cell type specificity peaks”) as a threshold to subset to high SAD score variants. For Enformer, we use the mean SAD scores across all chromatin accessibility tracks corresponding to the matched tissue (Blood/Immune tracks for Asthma and Eczema; Kidney tracks for Creatinine).

## Notes

### Competing Interest Statement

The authors have declared no competing interest.

